# Effective lowering of α-synuclein expression by targeting G-quadruplex structures within the SNCA gene

**DOI:** 10.1101/2024.02.09.579657

**Authors:** Valentina Pirota, Federica Rey, Letizia Esposito, Valentina Fantini, Cecilia Pandini, Erika Maghraby, Rosalinda Di Gerlando, Filippo Doria, Mariella Mella, Orietta Pansarasa, Paolo Gandellini, Mauro Freccero, Stephana Carelli, Cristina Cereda

**Affiliations:** Department of Chemistry, University of Pavia, Pavia, Italy; G4-INTERACT, USERN, Pavia, Italy; Pediatric Clinical Research Center “Fondazione Romeo ed Enrica Invernizzi”, Department of Biomedical and Clinical Sciences “L. Sacco”, University of Milan, Milan, Italy; Center of Functional Genomics and Rare diseases, Buzzi Children’s Hospital, Milan, Italy; Laboratory of Neurobiology and Neurogenetic, Golgi-Cenci Foundation, Abbiategrasso, Italy; Department of Biosciences, University of Milan, Milan, Italy; Department of Biology and Biotechnology “Lazzaro Spallanzani”, University of Pavia, Pavia, Italy; Molecular Biology and Transcriptomic Unit, IRCCS Mondino Foundation, Pavia, Italy; Cellular Models and Neuroepigenetics Unit, IRCCS Mondino Foundation, Pavia, Italy

**Keywords:** G-quadruplex, SNCA gene modulation, Alpha-synuclein, Parkinson’s disease

## Abstract

Alpha-synuclein, encoded by the SNCA gene, is a pivotal protein implicated in the pathogenesis of synucleinopathies, including Parkinson’s disease. Current approaches for modulating alpha-synuclein levels involve antisense nucleotides, siRNAs, and small molecules targeting SNCA’s 5’-UTR mRNA. Here, we propose a groundbreaking strategy targeting G-quadruplex structures to effectively modulate SNCA gene expression and lowering alpha-synuclein amount. Novel G-quadruplex sequences, identified on the SNCA gene’s transcription starting site and 5’-UTR of SNCA mRNAs, were experimentally confirmed for their stability through biophysical assays and *in vitro* experiments on human genomic DNA. Biological validation in differentiated SH-SY5Y cells revealed that well-known G-quadruplex ligands remarkably stabilized these structures, inducing the modulation of SNCA mRNAs expression, and the effective decrease in alpha-synuclein amount. Besides, a novel peptide nucleic acid conjugate, designed to selectively disrupt of G-quadruplex within the SNCA gene promoter, caused a promising lowering of both SNCA mRNA and alpha-synuclein protein. Altogether our findings highlight G-quadruplexes’ key role as intriguing biological targets in achieving a notable and successful reduction in alpha-synuclein expression, pointing to a novel approach against synucleinopathies.

## INTRODUCTION

Alpha-synuclein (α-Syn) is a presynaptic protein of 140 residues, encoded by the SNCA gene, widely expressed throughout the brain, with a high concentration in the presynaptic nerve terminals.[1] Although its physiological role is not yet fully characterized, it is mainly related to membrane binding, synaptic vesicle recycling, neurotransmitter release, and dopamine metabolism.[2] Under pathological conditions, impaired regulation of SNCA (including duplication or triplication of the SNCA gene locus) can induce an increase of α-Syn expression as high as 200% relative to its physiological levels. This leads to the accumulation of α-Syn monomers resulting in the formation of misfolded proteins that assemble into oligomers through the formation of β-sheet-rich fibrils.[3] These aggregates of α-Syn are the major components of Lewy Bodies, spherical filamentous protein inclusions that constitute the distinctive histopathological hallmarks in neurons of Parkinson’s disease (PD) affected brains.[3] The accumulation and release of α-Syn aggregates from affected brain regions can trigger synaptic and mitochondrial dysfunctions, changes in the lysosomal autophagy system, increased oxidative stress, as well as disruption of axonal transport, leading to neurodegeneration and progressive spreading of PD.[1,4]

In principle, these pieces of evidence suggest the potential of new therapeutic strategies aimed at lowering α-Syn production or increasing its clearance through the prevention and/or removal of its toxic aggregated deposits.[5] Many different approaches have been investigated in the last decade for preventing excessive buildup of α-Syn (a reduction by 25%–50% is expected to be sufficient to restore normal levels), and among them, the downregulation of SNCA gene expression attracted increasing interest in slowing or even stopping PD pathogenesis.[6] So far, promising results have been reached by epigenetic modification of SNCA through DNA methylation,[7] β2-adrenergic receptor agonists Clenbuterol and Salbutamol,[8] Gapmer-type antisense nucleotides,[9-11] short-interfering RNAs,[12] and small molecules acting on SNCA mRNAs.[13,14]

In this context, a new and promising strategy for lowering α-Syn production could be obtained by targeting G-quadruplex (G4) nucleic acid secondary structural motives. These are non-canonical high-order structures that occur in specific regions of the genome rich in guanines, like telomeres, gene promoters, transcription start sites (TSS), introns, and both 5’ and 3’ untranslated regions (UTRs).[15-18] Under physiological conditions, four guanine bases can self-associate via Hoogsteen-type hydrogen bonds base-pairing generating square-planar guanine quartets, known as G-tetrads. Two or more G-tetrads can stack on top of each other, through a cooperative stabilization due to monovalent cations (e.g. K^+^), constituting the backbone of G4 structures.[15] It is well-verified that G4s act as key structural elements in the regulation of a multitude of biological processes, among which is the modulation of gene expression.[19] Indeed, there are strong pieces of evidence that the stabilization of G4 structures in gene promoter regions interplays with the control of the transcriptional activity either increasing or decreasing it.[20,21] Moreover, G4s on 5’- and 3’-UTR mRNAs regions have been correlated with the regulation of the initial stage of the translation process, generally acting as structural silencing factors.[17] Therefore, the selective targeting of G4s, which has been reached successfully by small molecules,[22] is emerging as a novel therapeutic approach for a growing list of different pathologies, such as cancers,[16,18,23] bacterial and viral infections,[24,25] and neurodegenerative diseases.[26] About the last topic, it is important to note that G4s are often recognized as pathogenic drivers in various neurological disorders, with protective or deleterious effects in the cascade of neurotoxic processes.[26] Moreover, the well-defined folded nucleic acid structures of G4s make them advantageous therapeutic targets for upcoming drug development, mimicking the strategy of protein-targeted therapies.[21]

In this context, this work demonstrates, for the first time, the discovery and modulation of two new G4s, respectively located on the TSS of the SNCA gene (pSNCA) and on the 5’-UTR of the SNCA mRNA (mSNCA), in steering SNCA transcription process as well as in lowering α-Syn production. Biophysical studies demonstrated the folding of these sequences into very stable G4s under physiological conditions, which were further stabilized by well-known G4 ligands (G4Ls). In addition, the disruption of pSNCA G4 structure was achieved by a new conjugate based on a peptide nucleic acid sequence able to hybridize with pSNCA nucleo bases (NLS-PNA-C343). The treatment of differentiated SH-SY5Y cells with HPHAM[25,27] and Pyridostatin[28] G4-ligands, leads to the G4s’ stabilization, with an increase in the SNCA mRNAs expression and a simultaneous decrease in α-Syn protein levels. Conversely, the treatment with NLS-PNA-C343 leads to a decrease in both SNCA mRNAs and α-Syn expression.

Altogether these results emphasize the promising role of these new G4s as very promising targets for understanding SNCA regulation processes and reaching a novel and effective decrease in α-Syn production.

## RESULTS

### G-quadruplex folding prediction in SNCA promoter and transcripts

Putative quadruplex sequences (PQSs) were identified by using the online algorithm-based software “QGRS Mapper”,[29] “G4 Hunter”,[30] and “QuadBase2”,[31] where the likelihood of forming a stable G4 is evaluated by a G-score ranking. The SNCA gene is located on chromosome 4q22.1 (PARK1/4 loci), it contains 6 exons and spans about 114 kb. The main encoded transcript is SNCA140, translated by the last 5 exons of the gene. Other alternatively spliced transcripts encoding different isoforms have been identified for this gene, and in particular three (SNCA98, SNCA112, and SNCA126) result from alternative splicing of exons 3 and 5.[32] SNCA promoter region (Chr4: 89835200-89838801) is rich in G bases (22.4%), with a total %GC of 49.4% (Figure S1). Excluding the overlaps, 19 PQSs were identified, among which the only one with a QGRS Mapper G-score higher than 21 was pSNCA (G_4_ATG_4_CAG_5_CGCG_4_; location Chr4: 89837301-89837324; 24nt length) (Table S1). Moreover, this PQS was the only one also identified by “QuadBase2”[31] configuring medium stringency for G4 motives. pSNCA sequence is composed of four consecutive G-tracts of four guanines (Gs), respectively divided by a double-nucleotide tract (AT) and by two three-base sequences (CAG and CGC). It is located in the CpG island region, in the non-coding strand, and it straddles the TSS of the SNCA gene (-14/+10). Interestingly, this sequence shows QGRS Mapper and G4 Hunter score values higher than those of well-known G4s (Table 1). [33]

**Table 1.**
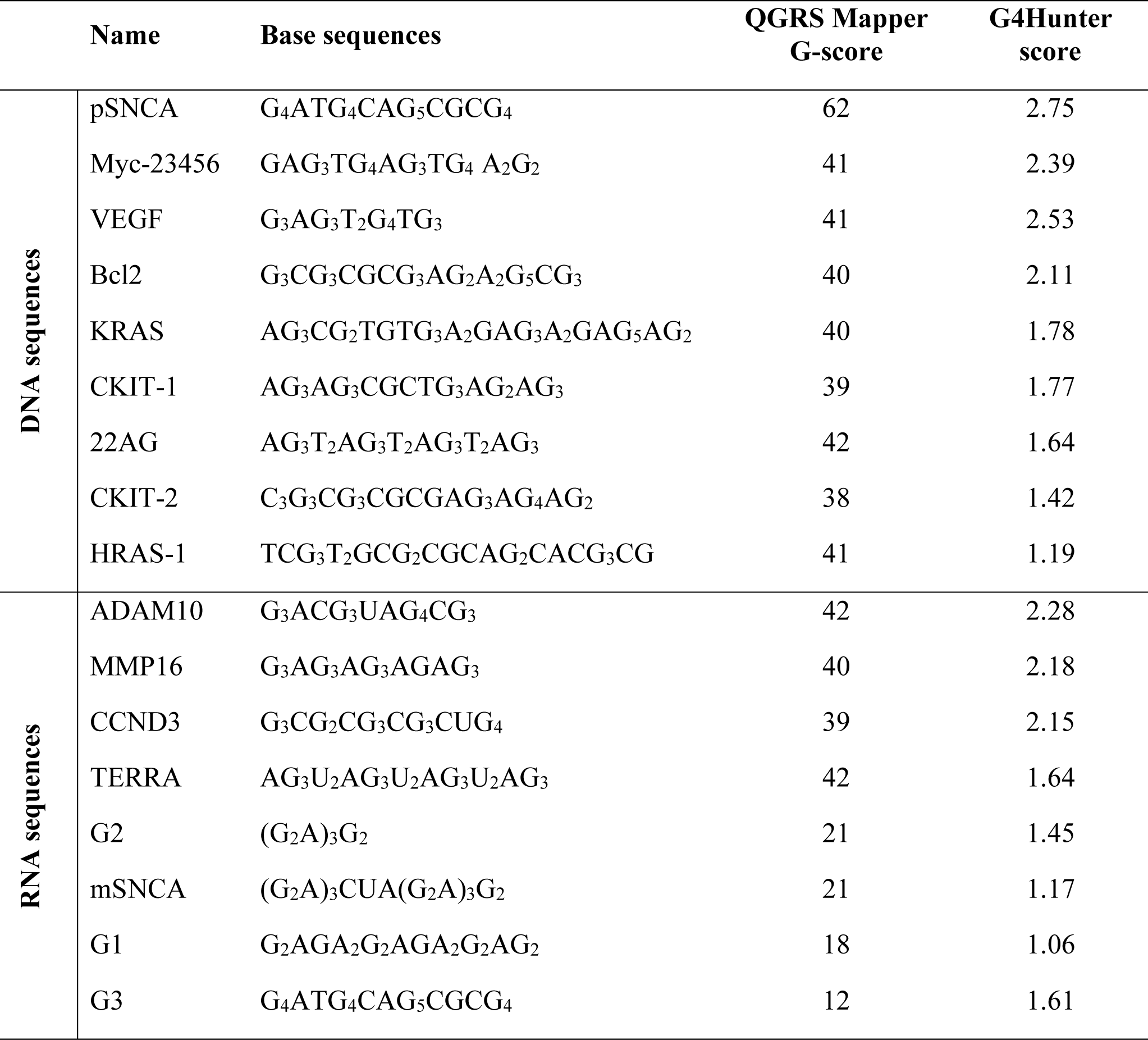
G-score values of pSNCA and mSNCA compared with the ones of well-known DNA and RNA G4s.

|  | Name | Base sequences | QGRS Mapper<br>G-score | G4Hunter<br>score |
| --- | --- | --- | --- | --- |
| DNA sequences | pSNCA | G<sub>4</sub>ATG<sub>4</sub>CAG<sub>5</sub>CGCG<sub>4</sub> | 62 | 2.75 |
|  | Myc-23456 | GAG<sub>3</sub>TG<sub>4</sub>AG<sub>3</sub>TG<sub>4</sub> A<sub>2</sub>G<sub>2</sub> | 41 | 2.39 |
|  | VEGF | G<sub>3</sub>AG<sub>3</sub>T<sub>2</sub>G<sub>4</sub>TG<sub>3</sub> | 41 | 2.53 |
|  | Bcl2 | G<sub>3</sub>CG<sub>3</sub>CGCG<sub>3</sub>AG<sub>2</sub>A<sub>2</sub>G<sub>5</sub>CG<sub>3</sub> | 40 | 2.11 |
|  | KRAS | AG<sub>3</sub>CG<sub>2</sub>TGTG<sub>3</sub>A<sub>2</sub>GAG<sub>3</sub>A<sub>2</sub>GAG<sub>5</sub>AG<sub>2</sub> | 40 | 1.78 |
|  | CKIT-1 | AG<sub>3</sub>AG<sub>3</sub>CGCTG<sub>3</sub>AG<sub>2</sub>AG<sub>3</sub> | 39 | 1.77 |
|  | 22AG | AG<sub>3</sub>T<sub>2</sub>AG<sub>3</sub>T<sub>2</sub>AG<sub>3</sub>T<sub>2</sub>AG<sub>3</sub> | 42 | 1.64 |
|  | CKIT-2 | C<sub>3</sub>G<sub>3</sub>CG<sub>3</sub>CGCGAG<sub>3</sub>AG<sub>4</sub>AG<sub>2</sub> | 38 | 1.42 |
|  | HRAS-1 | TCG<sub>3</sub>T<sub>2</sub>GCG<sub>2</sub>CGCAG<sub>2</sub>CACG<sub>3</sub>CG | 41 | 1.19 |
| RNA sequences | ADAM10 | G<sub>3</sub>ACG<sub>3</sub>UAG<sub>4</sub>CG<sub>3</sub> | 42 | 2.28 |
|  | MMP16 | G<sub>3</sub>AG<sub>3</sub>AG<sub>3</sub>AGAG<sub>3</sub> | 40 | 2.18 |
|  | CCND3 | G<sub>3</sub>CG<sub>2</sub>CG<sub>3</sub>CG<sub>3</sub>CUG<sub>4</sub> | 39 | 2.15 |
|  | TERRA | AG<sub>3</sub>U<sub>2</sub>AG<sub>3</sub>U<sub>2</sub>AG<sub>3</sub>U<sub>2</sub>AG<sub>3</sub> | 42 | 1.64 |
|  | G2 | (G<sub>2</sub>A)<sub>3</sub>G<sub>2</sub> | 21 | 1.45 |
|  | mSNCA | (G<sub>2</sub>A)<sub>3</sub>CUA(G<sub>2</sub>A)<sub>3</sub>G<sub>2</sub> | 21 | 1.17 |
|  | G1 | G<sub>2</sub>AGA<sub>2</sub>G<sub>2</sub>AGA<sub>2</sub>G<sub>2</sub>AG<sub>2</sub> | 18 | 1.06 |
|  | G3 | G<sub>4</sub>ATG<sub>4</sub>CAG<sub>5</sub>CGCG<sub>4</sub> | 12 | 1.61 |

The 5′-UTR of the human SNCA mRNA is a 264 nt long G-rich (41.2%) region, with an overall 65.5% of GC content. Previous investigation of this region led to the discovery of three non-overlapping PQS motifs (G1, G2, and G3 in Table 1)[34] with rather subpar QGRS Mapper and G4 Hunter scores compared to most verified mRNA G4 structures (Table 1). Nevertheless, considering that their combined effect resulted in 45% translation inhibition,[34] the 5′-UTR of the human SNCA mRNA was further analyzed, leading to the discovery of a new PQS that spans G1 and G2 sequences. This novel sequence, called mSNCA (Table 1; GA_2_(G_2_A)_3_CUA(G_2_A)_3_G_2_; 23 nt length), contains two repetitions of the GGA triplet belonging to G1, a GGACUA hexanucleotide stretch, and three repetitions of the GGA triplet with a terminal two-bases tract (GG) belonging to G2.

### Confirmation of G4s folding in SNCA promoter and transcripts

#### Chromatin Immunoprecipitation and qPCR stop assay

To verify the pSNCA G4-folding at the cellular setting, Chromatin Immunoprecipitation (ChIP) and quantitative PCR (qPCR) stop assays were performed. Both ChIP and qPCR stop assays were accomplished on genomic DNA from SH-SY5Y cells after 7 days of neural differentiation.

Primers covering an 182 base-pair region (SNCA_G4-FW and SNCA_G4-REV) centered on TSS were used to check the folding of pSNCA into a G4 structure by G4-ChIP. This assay showed the enrichment of SNCA-G4s DNA upon chromatin immunoprecipitation with the anti-G4 antibody BG4[35] as compared to an anti-FLAG antibody (Figure 1A). In contrast, a non-G4 forming GAPDH region, used as a negative control, was not enriched, emphasizing that the pSNCA sequence folds into a G4 structure *in vitro*. qPCR stop assay was performed by applying different potassium chloride concentrations (from 0 to 50 mM), exploiting the same SNCA-G4 primers used for G4-ChIP. Stabilization of SNCA G4 structure by K^+^ ions caused an obstacle for the DNA polymerase during DNA amplification, with a reduction of amplicons even with 0.1 mM KCl (Figure 1B). This trend parallels the results obtained for MYC, which was used as a positive control, highlighting the high stability of this new SNCA-G4 (Figure S2). Primers spanning a non-G4 sequence in the GAPDH gene were used as a negative control, and indeed its amplification products did not show a significant alteration in response to the increase in K^+^ (Figure S2). Additionally, the presence of G4s in the 5’-UTR-RNA region of SNCA was demonstrated by G4-RIP analysis highlighting a specific enrichment for the mRNA SNCA-G4 (Figure 1C). The qPCR stop assay performed on cDNA also showed a slightly decreased target amplification at increasing concentrations of KCl (Figure 1D).

**Figure 1.**
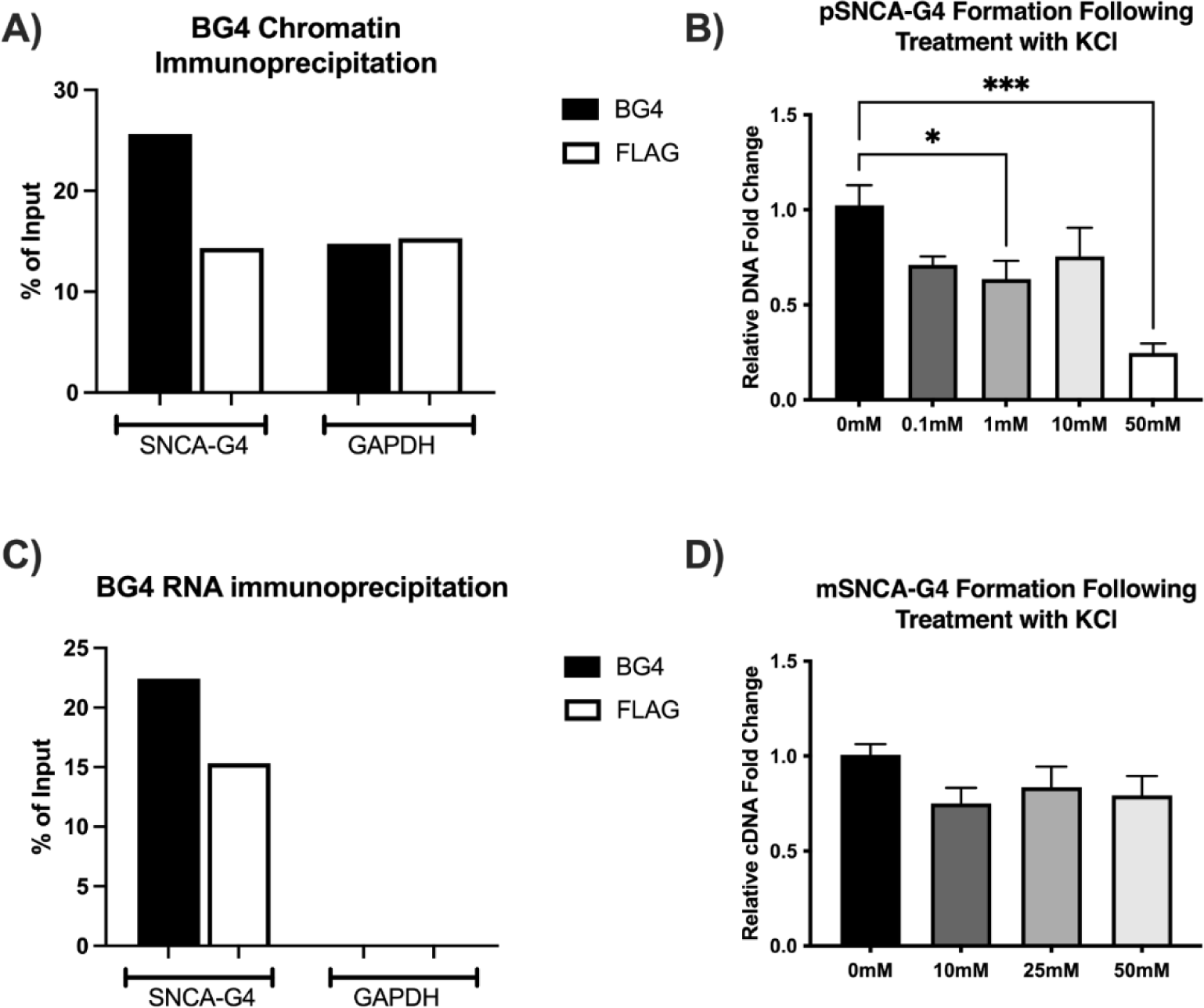
Cellular validation of the presence of G4 structures within SNCA promoter and SNCA 5’-UTR mRNA. A) BG4 and anti-FLAG antibodies were used for chromatin immunoprecipitation and enrichment for the SNCA-G4 DNA region was determined by qPCR. A non-G4 forming GAPDH region was used as a negative control. B) SNCA-G4 abundance at promoter level was assayed via qPCR stop assay with increasing KCl concentrations, using a non-G4 forming GAPDH region as a housekeeping gene. Data are expressed as mean ± SEM of 5 independent samples (n = 5, *p<0.05, ***p<0.001 vs 0mM). C) BG4 and FLAG were used for RNA immunoprecipitation and enrichment for the SNCA-G4 cDNA region was determined by qPCR. A non-G4 forming GAPDH region was used as a negative control. D) SNCA-G4 abundance at 5’-UTR mRNA level was assayed via qPCR stop assay with increasing potassium chloride concentrations, using a non-G4 forming GAPDH region as a housekeeping gene. Data are expressed as mean ± SEM of 5 independent samples (n = 6). All assays were performed on SH-SY5Y cells after 7 days of neural differentiation.

#### CD, UV-vis, FRET, and NMR Studies on SNCA model oligonucleotides

Biophysical investigations were carried out to confirm that these two sequences can fold into G4 structures. The capacity of the pSNCA DNA sequence (G_4_ATG_4_CAG_5_CGCG_4_, 24nt) to fold into G4 was confirmed using circular dichroism (CD), UV-vis thermal difference spectra (TDS), and preliminary one-dimensional ^1^H-NMR. CD spectra were recorded in the presence of different KCl concentrations (from 0 to 100 mM), showing a mainly antiparallel topology (calculated equal to 64%, Figure S3A),[36] with a positive peak around 295 nm and a negative one around 260 nm (Figure 2A), in the presence of high potassium concentrations (100 mM K^+^).[37] As the potassium concentration decreases, the sequence rebalances itself towards a 3+1 hybrid topology, highlighted by the growth of a positive shoulder at around 270 nm (hybrid topology percentage equal to 68% with 0.5 mM K^+^, Figure S3A). To strengthen the biophysical model making it more consistent with the real biological SNCA G4-folding equilibria, an extended sequence, pSNCAext (GATG_4_ATG_4_CAG_5_CGCG_4_TGA, 30-nt), that includes the 3 previous nucleobases (GAT) and the 3 subsequent nucleobases (TGA) present at the genomic level, was also studied. In this case, the addition of 6 nucleobases, including two more guanines, shifts the equilibrium towards a 3+1 hybrid topology (92% with 100 mM K^+^, Figure S3C), characterized by a positive peak at around 295 nm with a shoulder at around 275 nm (Figure 2B).[37] By decreasing potassium chloride, a transition to a mixture of hybrid and antiparallel topologies (equal to 62.1% and 37.9% respectively with 0.5 mM K^+^, Figure S3C) was obtained. A complete analysis of estimated secondary and tertiary structure fractions of pSNCA and pSNCAext G4s as a function of potassium chloride concentration is reported as Supporting Information (Figure S3). TDS spectra, obtained by subtracting the spectra of unfolded (temperature above T_m_) and folded (low temperature) G4s, revealed G4’s characteristic peaks: a negative one at 295 nm, and positive ones at 273 nm, 257 nm, and 243 nm (Figures 2C-D).[38] The computed TDS factors (ΔA240nm/ΔA295nm), equal to 1.06 ± 0.08 for pSNCA and 1.3 ± 0.1 for pSNCAext supported their folding into antiparallel (group III) and hybrid (group II) topologies.[38] Further confirmation came from ^1^H-NMR signals in the 10-12 ppm range (Figures 2E-F), which are typical of Gs imino protons when a G-tetrad is formed, as opposed to 13-14 ppm for those involved in Watson-Crick base pairing.[39] CD-melting experiments on both the model oligonucleotides described their folding into extremely stable G4s, showing mid-transition temperature (T_m_) values equal to 61.7 ± 0.1°C and 56.8 ± 0.1°C for pSNCA and pSNCAext respectively in the presence of only 1 mM K^+^. This potassium chloride concentration is over two orders of magnitude lower than the physiological value in the cell (around 150 mM),[40] but the high stability of these two new G4s makes it difficult to have a correct measurement of T_m_ values by biophysical methods already at 50 mM KCl. Indeed, by CD-melting, T_m_ values are very close to the detection limit set at 95°C with 50 mM K^+^ (94.3 ± 0.7°C and 91.8 ± 0.3 °C respectively for pSNCA and pSNCAext) and over 95°C with 100 mM K^+^. To further underline the remarkable stability of these pSNCA-G4s, we performed CD-melting experiments on well-known G4-models (Myc-23456, cMyc, 22ag, CKIT-1, and LTR-III) [33,41] with different KCl concentrations to maintain T_m_ values in the 50-60 °C range. It should be noted that the model sequence of the most stable parallel G4 known, Myc-23456, showed a T_m_ of 61.2 ± 0.4°C in the presence of 5 mM K^+^, about 12 degrees lower than T_m_ obtained with pSNCA (74.6 ± 0.1°C) and pSNCAext (73.2 ± 0.6°C) in the same conditions (Table S2). Following the absorbance decrease at 295 nm, mid-transition temperature values for pSNCA and pSNCAext were determined to be 61.5 ± 0.4°C and 55.1 ± 0.1 °C, respectively,[42] excellent agreement with CD-melting data. From the UV-melting curves, model-dependent thermodynamic parameters were extracted to better evaluate the thermodynamics of these novel G4s (Table S3).[42] Fluorescence resonance energy transfer (FRET)-melting experiments, using 5’-FAM/3’-TAMRA labeled oligonucleotides, corroborated mid-transition temperature data. Following the G4-unfolding of 0.25 µM model oligonucleotides through a temperature-dependent increase in donor emission of FAM (25-95°C temperature range) in the presence of 1 mM KCl, we measured pSNCA and pSNCAext T_m_ equal to 64.3 ± 0.4°C and 58.8 ± 0.8 °C, respectively.[43] Supporting Information contains a summary of the computed stabilities as a function of potassium chloride concentrations, determined by exploiting the thermal unfolding of pSNCA and pSNCAext G4s by different techniques (Table S4).

**Figure 2.**
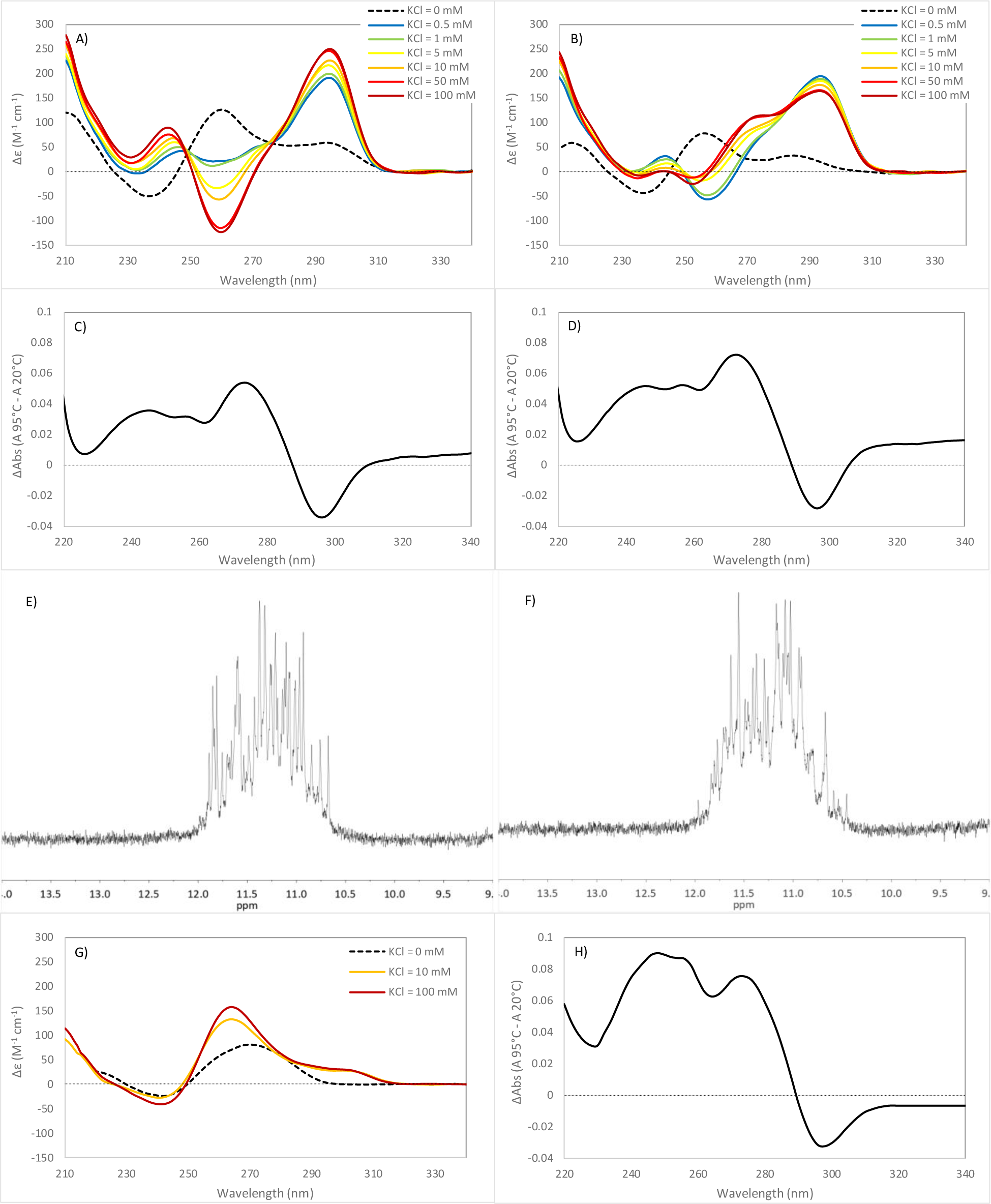
Biophysical data supporting the folding of SNCA model oligonucleotides into G4. CD spectra of 2.5 µM of A) pSNCA and B) pSNCAext in 10 mM lithium cacodylate buffer (pH 7.4) with different potassium concentrations (ranging from 0 to 100 mM). The ionic strength is maintained constant at 100 mM by adding proper amounts of LiCl. TDS spectra of 2.5 µM C) pSNCA and D) pSNCAext in 10mM lithium cacodylate buffer pH 7.4, with 1 mM KCl and 99 mM LiCl. ^1^H-NMR of 0.25 mM E) pSNCA and F) pSNCAext in 90% H_2_O/10% D_2_O solution with 100 mM KCl and 20 mM phosphate buffer pH 7.4, at 25°C. G) CD spectra of 2.5 µM of mSNCA in 10 mM lithium cacodylate buffer (pH 7.4) with different potassium concentrations (0-10-100 mM). The ionic strength is maintained constant at 100 mM by adding proper amounts of LiCl. H) TDS spectra of 2.5 µM mSNCA in 10 mM lithium cacodylate buffer pH 7.4, with 100 mM KCl.

As no biophysical analysis was previously conducted on the PQSs identified in the 5’-UTR-RNA region of SNCA, a deep biophysical analysis was performed on mSNCA model oligonucleotide ((G_2_A)_3_CUA(G_2_A)_3_G_2_, 23 nt) to confirm its propensity to fold into a stable mRNA G4.

Dichroic spectra of mSNCA (2.5 µM) recorded at different potassium concentrations (0, 10, 100 mM) in 10 mM lithium cacodylate buffer pH 7.4, confirmed its folding into a parallel G4 structure, characterized by a positive peak at 264 nm and a negative one at around 240 nm (Figure 2G). The very small positive peak observed at 300-310 nm may be associated with the stacking of mixed G/A heptads and/or hexads, structural motifs recognized to develop in RNA/DNA sequences that feature repetitions of GGA.[44] Additionally, mSNCA folding into a G4 structure was confirmed by TDS, which revealed a negative peak at around 295 nm, and positive ones at around 273 nm, 257 nm, and 243 nm (Figure 2H).[38] The calculated TDS factor (ΔA240nm/ΔA295nm) of 2.4 ± 0.2 highlighted its folding into a mainly parallel G4 (group I).[38]

CD and UV melting experiments in 10 mM lithium cacodylate buffer pH 7.4 and in the presence of 100 mM KCl, showed a melting temperature of 62.4 ± 0.1 °C, as well as FRET-melting analysis revealed a T_m_ = 68.5 ± 0.6 in the same experimental conditions. These results emphasized that mSNCA is a pretty stable mRNA G4 when compared to other well-known mRNA G4s (TERRA, ADAM10, MMP16)[23,45,46] which melt at around 60°C in the presence of much less potassium chloride (Table S2), following the G-score values identified by bioinformatic analysis. A summary of melting data was reported in Table S4. Model-dependent thermodynamic parameters were extracted from the UV-melting curve (Table S3).[42]

In addition, a Principal Component Analysis (PCA) on a reference library of 23 known G4s CD spectra was conducted to highlight pSNCA, pSNCAext, and mSNCA topologies features.[36] The scores plot of the first two components PC1 (explained variance 62.82%) and PC2 (explained variance 18.87%) showed the separation of three clusters, corresponding to parallel, hybrid, and antiparallel G4 topologies (Figure S4). The CD spectrum of pSNCAext perfectly fits into the G4s set relating to the hybrid topology, as well as the CD spectrum of pSNCA fits within the antiparallel G4s group and that of mSNCA in the parallel one.

### SNCA-G4 folding modulation by well-known G4-ligands and PNA conjugates

#### FRET- and CD-melting experiments to select the best-stabilizing G4 ligand

To define the involvement of G4s in the modulation of SNCA gene expression, pSNCA and pSNCAext stabilization were measured in the presence of four well-known G4 ligands. Among the countless known G4 binders, we selected the well-known HPHAM (core-extended naphthalene diimide), PDS (Pyridostatin, a bis-quinoline derivative), BRACO19 (acridine-based ligand), and TMPyP4 (porphyrin-based ligand).[22]

From FRET- and CD-melting results, the three ligands HPHAM, PDS, and BRACO19 stabilize to different extents both model sequences, with the best-performing stabilizer being HPHAM, which is able to stabilize both model oligonucleotides of around 13-16°C (Table 2). On the contrary, TMPyP4 destabilized both the model structures, from 2 to 11°C, consistent with its sporadically described propensity to unfold some G4 structures.[47] In addition, when analyzing the effect of the binding on the native topology of SNCA-G4s, we realized that among the analyzed binders, only HPHAM can stabilize both pSNCA and pSNCAext G4 structures preserving their original topology unaltered (Figure S5). This is confirmed at both 1 mM and 100 mM KCl concentrations, corresponding respectively to the working concentration of FRET-/CD-melting and an experimental environment with a physiological-like potassium ions concentration.

**Table 2.**
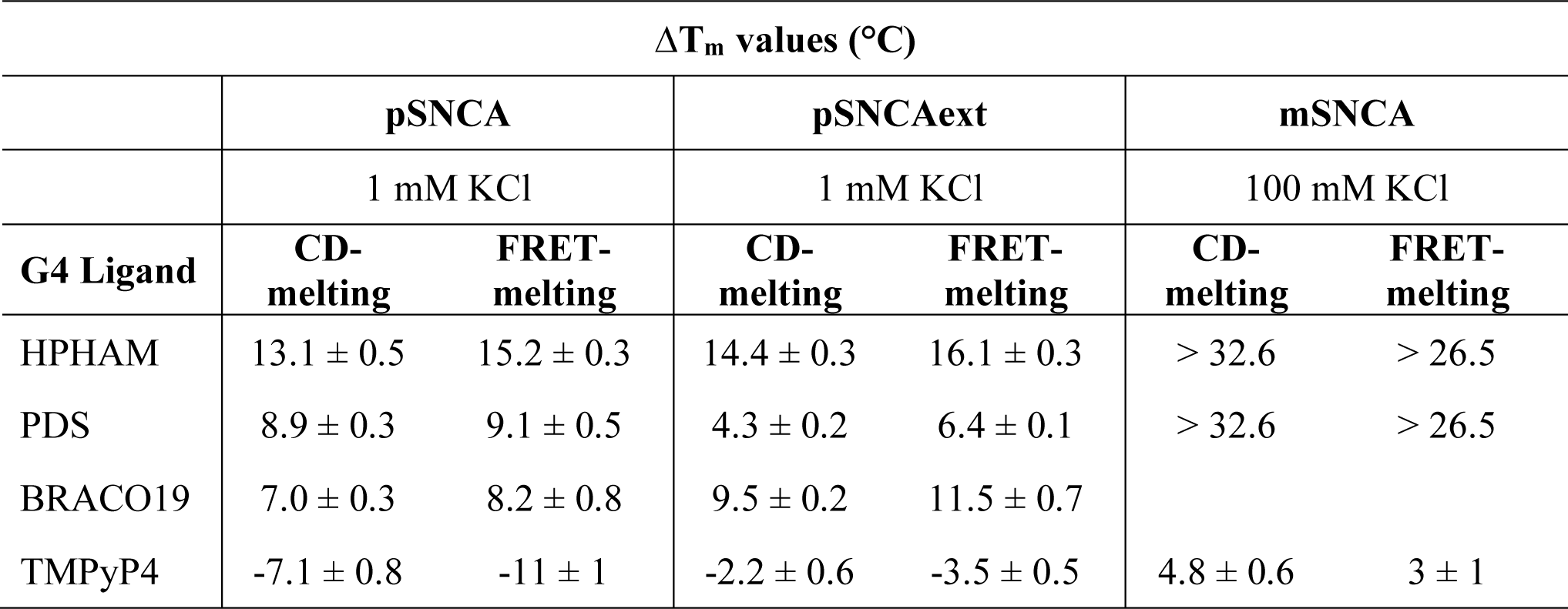
Melting data summary describing the G4-stabilization/destabilization induced by G4-ligands. pSNCA (or pSNCAext, or mSNCA) 2.5µM and 0.25µM for CD and FRET melting experiments respectively, in the presence of 4 equiv. of G4-ligands, and 10 mM lithium cacodylate buffer pH 7.4. Data for pSNCA and pSNCAext were recorded with 1 mM KCl and 99 mM LiCl. Data for mSNCA were recorded with 100 mM KCl.

| $\Delta T_m$ values (°C) | | | | | | |
| --- | --- | --- | --- | --- | --- | --- |
|  | pSNCA |  | pSNCAext |  | mSNCA |  |
|  | 1 mM KCl |  | 1 mM KCl |  | 100 mM KCl |  |
| G4 Ligand | CD-melting | FRET-melting | CD-melting | FRET-melting | CD-melting | FRET-melting |
| HPHAM | 13.1 ± 0.5 | 15.2 ± 0.3 | 14.4 ± 0.3 | 16.1 ± 0.3 | > 32.6 | > 26.5 |
| PDS | 8.9 ± 0.3 | 9.1 ± 0.5 | 4.3 ± 0.2 | 6.4 ± 0.1 | > 32.6 | > 26.5 |
| BRACO19 | 7.0 ± 0.3 | 8.2 ± 0.8 | 9.5 ± 0.2 | 11.5 ± 0.7 |  |  |
| TMPyP4 | -7.1 ± 0.8 | -11 ± 1 | -2.2 ± 0.6 | -3.5 ± 0.5 | 4.8 ± 0.6 | 3 ± 1 |

Considering the excellent results obtained with HPHAM, its stabilizing power was assessed on the mSNCA model oligonucleotide. PDS was also tested as the best-known binder reference, as well as TMPyP4 for its destabilizing effect on pSNCA. In the presence of 100 mM KCl, both PDS and HPHAM strongly stabilized the mSNCA G4 structure so much so that the melting temperature could not be determined within the 95 °C (Table 2). Otherwise, TMPyP4 only slightly stabilized the mSNCA structure, increasing the melting temperature to no more than 5 °C (Table 2).

#### CD study to test the ability of a new PNA conjugate to disrupt pSNCA G4 structure

In parallel with the identification of good binders capable of strongly stabilizing its structure, a short peptide nucleic acids (PNA)[48] sequence (C_4_GTC_3_) capable of hybridizing with the central bases of the pSNCA (or pSNCAext, from G10 to G18, Figure S6) was synthesized. A nuclear localization sequence (NLS) H_2_N-ProLysLysLysArgLysVal-OH was conjugated to the PNA C-terminus, with the inclusion of a glycine residue as a spacer, to accomplish PNAs delivery into the nuclei. In addition, a Coumarin 343 (C343) residue was coupled to the N-terminus of PNA by separating the dye through a lysine residue, which is positively charged and increases the electrostatic contact between the two strands, implementing water-solubility and cellular uptake. The synthesis of the new NLS-PNA-C343 conjugate (H_2_NCO-NLS-Gly-C_4_GTC_3_-Lys-C343) was accomplished using Fmoc solid-phase peptide synthesis (SPPS) methodology, by exploiting a rink amide resin with a low loading (0.37 mmol/g) to enhance synthetic efficiency. Coumarin 343 was added as the last synthesis step by coupling reaction in DMF (r.T., 2h). In parallel, an NLS-PNA5 sequence (H_2_NCO-NLS-Gly-C_2_GCAC_2_G_2_-Lys) with five mismatched bases to pSNCA (Figure S6) was synthesized to verify the PNA strategy’s sensitivity on target recognition.[49] The power of the two new PNA-conjugates was tested in the interaction with the model sequence pSNCAext to better mimic their effect on an extended region. pSNCAext (2.5 µM) was folded into the corresponding G4 structure in the presence of 100 mM KCl in 10 mM lithium cacodylate buffer, pH 7.4. Subsequently, 4 equivalents of NLS-PNA-C343 or NLS-PNA5 were added and the solution was left to equilibrate for 4 hours at 25°C before recording the CD spectra. Although a PNA/DNA heteroduplex typically forms a 1:1 complex, 4 equivalents of PNA conjugates were employed to ensure effective competition with the highly stabilizing nature of pSNCAext’s G4 structure and to enable a more accurate comparison of PNAs’ contribution relative to stabilizing ligands (studied at the same molar concentration). In the presence of NLS-PNA-C343, a drastic change in dichroic signal with a positive band around 278 nm and a negative one around 252 nm was obtained, confirming the PNA-DNA duplex formation, following G4 unfolding (Figure S6). The formation of a B-form structure was confirmed by analyzing the CD spectrum with the CD-NuSS web server using the XGBoost algorithm prediction method (accuracy 87.33%).[50] The melting temperature of this new hetero duplex was evaluated by CD-melting and resulted in 54.3 ± 0.5 °C. Otherwise, the same experiment with NLS-PNA5 resulted in a modest blue shift of the pSNCAext CD spectrum, which remains primarily a hybrid G4. The unfolding temperature for the small percentage of PNA-DNA heteroduplex present was 42.5 ± 0.5 °C (Figure S6). Furthermore, CD spectra data was incorporated as a new sample in the conducted PCA to highlight the significant disparity in the dichroic spectra features. This analysis corroborates the observation that the dichroic characteristics of the new structure deriving from pSNCAext interaction with NLS-PNA-C343 do not align with those of established G4s, lacking the requisite features to fit within any of the three PCA clusters (Figure S4). Differently, the spectra observed in the presence of NLS-PNA5 remained in the G4 set corresponding to the hybrid topology, indicating that it did not significantly affect the G4 structure of pSNCAext (Figure S4).

### Evaluation of SNCA mRNA and α-Syn protein expression following G4 stabilization

To assess the impact of SNCA-G4 stabilization on SNCA mRNA and α-Syn expression, *in vitro* experimental assays on the SH-SY5Y cell line following neural differentiation were performed.[51,52] Considering the biophysical results, we selected HPHAM and PDS as the best stabilizing G4-ligands to test, as well as TMPyP4 for its destabilizing activity on pSNCA G4. Their non-cytotoxic concentrations were first identified in the SH-SY5Y cell line by MTT assay, defying (i) 625 nM as the highest non-toxic dosage for HPHAM, even after 48 hours of treatment (Figure S7); (ii) 5 µM for PDS, in complete agreement with already reported studies on the SH-SY5Y cell line (Figure S8);[53,54] and (iii) 8 µM for TMPyP4 (Figure S9). In addition, taking advantage of the intrinsic red fluorescence emission of HPHAM, we highlighted that HPHAM efficiently enters cells at 625 nM, already after 4 hours (Figure S10). By Real-Time PCR we identified an increase over time in SNCA mRNA expression following HPHAM treatment (Figure 3A), whilst it remained essentially stable for up to 48 hours when cells were treated with PDS (Figure 3B) and TMPyP4 (Figure S11A). The G4-ligands activity was validated by assessing the expression of MYC as a positive control, which, as expected, was reduced following G4-ligand treatments (Figures 3A, 3B, and S11B). TATA Binding Protein (TBP) was used as a housekeeping gene as we found this as the most stable housekeeping not influenced by G4 stabilization (as opposed to GAPDH and 18S; Figure S12). To evaluate whether SNCA was newly synthesized, a decay assay was performed treating cells with both HPHAM or PDS and the transcriptional inhibitor Actinomycin D (Figures 3C and 3D).[55] In both cases, following treatment with the G4-stabilizer, a faster decay of RNA expression was emphasized, most significant after 8 hours of treatment (around 50% with both ligands).

**Figure 3.**
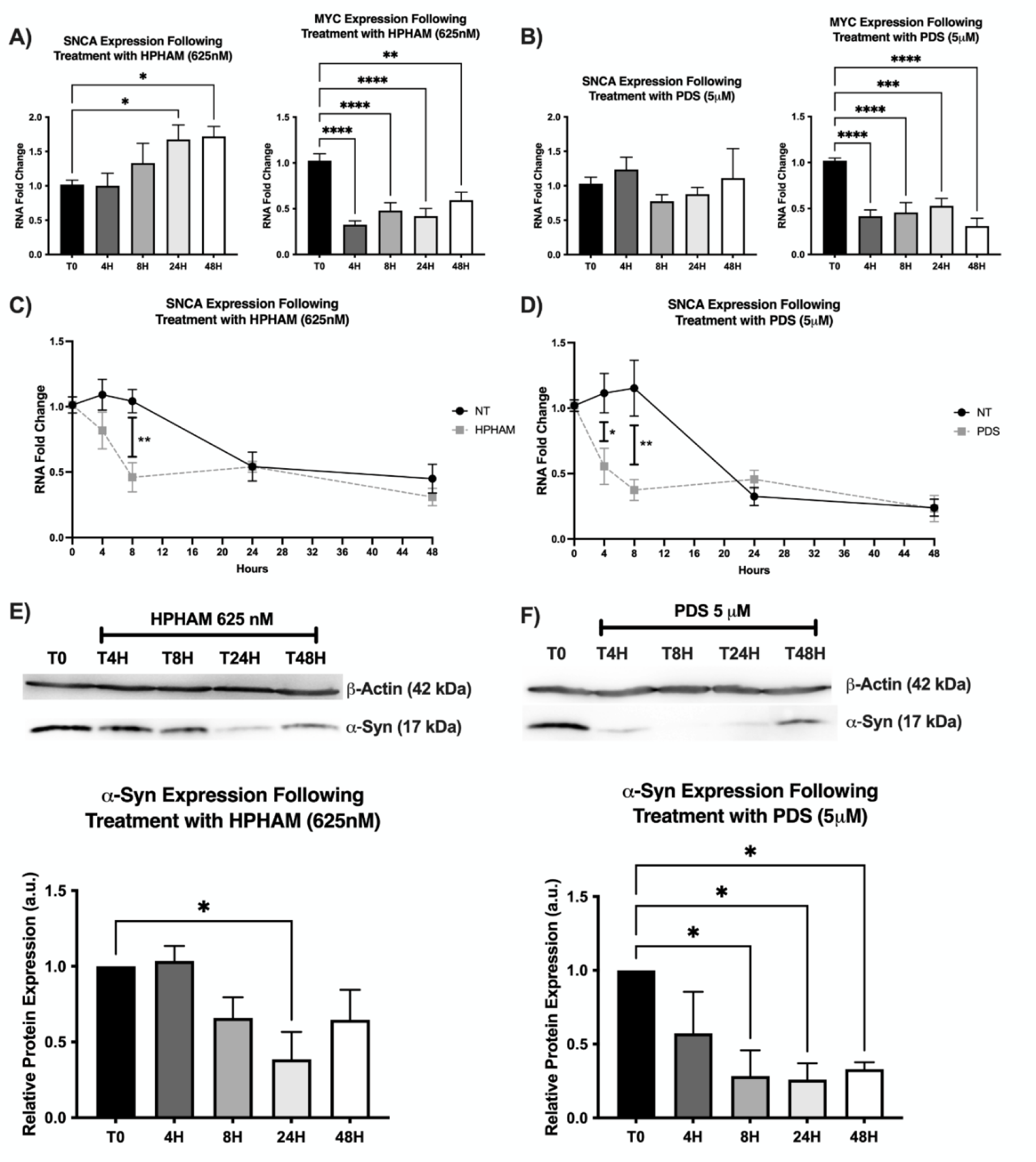
Assessment of SNCA mRNA and α-Syn expression as a consequence of G4 stabilization. SNCA and MYC mRNA expression levels were assessed via Real-Time PCR following treatment with 625 nM HPHAM (A) and 5 µM PDS (B) for 0, 4, 8, 24, and 48 hours. TBP was used as a housekeeping gene and data are expressed as mean ± SEM of 3 replicate values in 3 independent experiments (n=9; *p<0.05, **p<0.01, ***p<0.001, ****p<0.0001 vs T0). SNCA expression levels were also assessed via Real-Time PCR following treatment with 1.5 μg/mL Actinomycin D and either 625 nM HPHAM (C) or 5 µM PDS (D) for 0, 4, 8, 24, and 48 hours. α-Syn expression levels were assessed via Western Blotting following treatment with 625 nM HPHAM (E) and 5 µM PDS (F) for 0, 4, 8, 24, and 48 hours. β-Actin was used as a housekeeping protein and data are expressed as mean ± SEM of 3 independent experiments (n=3; *p<0.05 vs T0).

To assess the impact of G4-stabilizing ligands on α-Syn protein expression, Western Blotting analyses were performed on protein extracts obtained from differentiated SH-SY5Y cells treated with HPHAM, PDS, or TMPyP4 at previously used non-cytotoxic concentrations (625 nM, 5 µM, and 8 µM respectively) for 0, 4, 8, 24, and 48 hours. In this case, β-actin was used as a housekeeping protein as its expression is not impacted by treatment with the G4-binders. HPHAM and PDS administration resulted in a significant reduction in protein expression, with α-Syn protein drop by more than 50% after 24h (Figures 3E-3F). In contrast, the treatment with TMPyP4 did not affect α-Syn protein expression for 24 hours, but a slight increase was observed after 48 hours (Figure S13).

### Selective disruption of TSS G4 leads to the reduction in SNCA mRNA and α-Syn expression

To better understand the specific role of the G4 structure within the promoter region of the SNCA gene, NLS-PNA-C343 was tested for its ability to disrupt the pSNCA G4 folding. NLS-PNA-C343 resulted in being non-toxic at 5 µM dosage after 24 hours (Figure S14), thus this dosage was selected for further experiments. Following 7 days of neural differentiation, cells were treated with NLS-PNA-C343 for 0, 4, 8, and 24 hours (5 µM). The effective disruption of pSNCA G4 drove a decrease in SNCA mRNA expression of around 50% after only 4 hours (Figure 4A), conversely to results obtained with G4s’ stabilizing ligands. Moreover, this selective targeting, transitioning from an initial enhancement of α-Syn production after 4 hours, culminated in a substantial decrease in α-Syn expression, reaching approximately 70% reduction after 24 hours (Figure 4B). The five-bases mismatched NLS-PNA5 was tested at 5 µM dosage for comparison (Figure S15A), demonstrating the effectiveness of the antisense technique in ensuring high target selectivity. Indeed, neither mRNA levels (Figure S15B) nor α-Syn expression were significantly impacted by the mismatched NLS-PNA5 treatment (Figure S15C).

**Figure 4.**
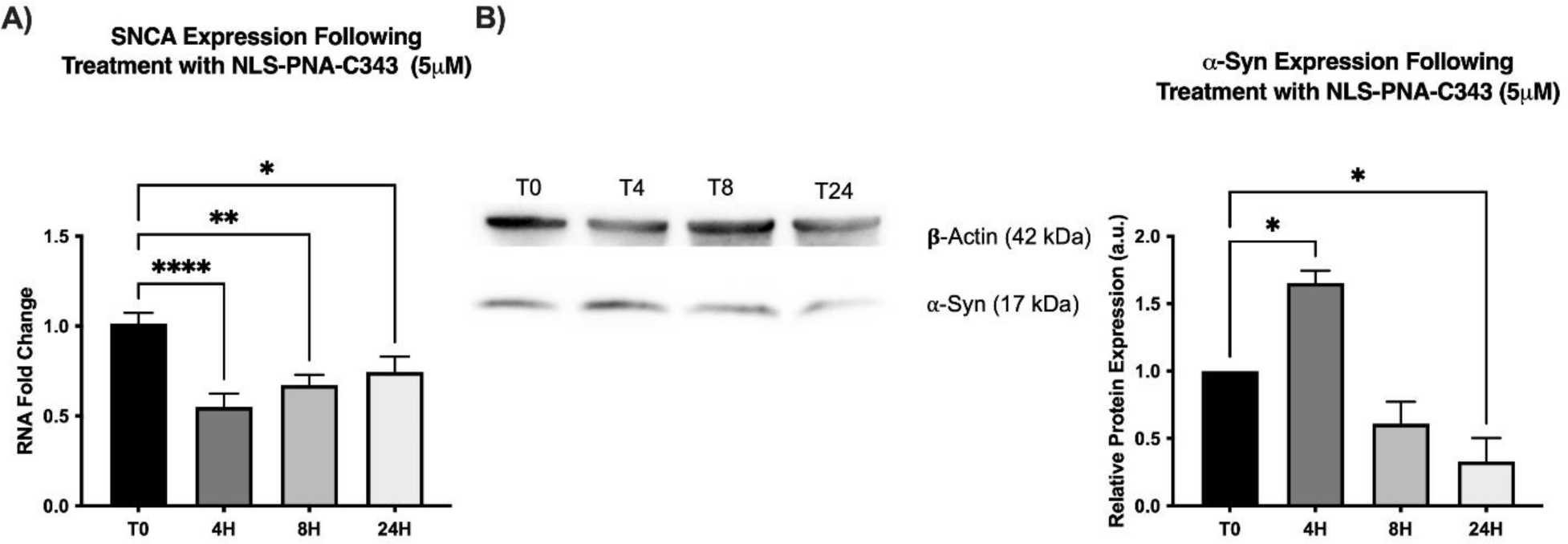
Assessment of SNCA mRNA and α-Syn expression as a consequence of selective pSNCA G4 destabilization. (A) SNCA mRNA expression levels were assessed via Real-Time PCR following treatment with 5 µM SNCA-G4 destabilizer for 0, 4, 8, and 24 hours. TBP was used as a housekeeping gene and data are expressed as mean ± SEM of 3 replicate values in 3 independent experiments (n=9; *p<0.05, **p<0.01, ****p<0.0001 vs T0). (B) α-Syn expression levels were assessed via Western Blotting following treatment with 5 µM SNCA-G4 destabilizer for 0, 4, 8, and 24 hours. β-Actin was used as a housekeeping protein and data are expressed as mean ± SEM of 2 independent experiments (n=2; *p<0.05 vs T0).

## DISCUSSION

Dysregulation of α-Syn production in pathologic conditions leads to protein aggregates linked to Parkinson’s disease and other synucleinopathies, highlighting the need for therapeutic strategies aimed at reducing α-Syn synthesis.[5] In this context, targeting secondary nucleic acid structures, such as G-quadruplexes, to regulate transcription and translation is a novel and high-impact challenge.[19] Here, the potential to target G4 structures to modulate α-Syn expression has been highlighted through bioinformatic identification of two promising PQSs on the SNCA gene. One of these sequences spans the TSS of the SNCA gene (pSNCA), and the other is located within the 5’-UTR of SNCA transcripts in the internal ribosome entry site (mSNCA). Specifically, the location of pSNCA within the TSS of the SNCA gene (-14/+10) suggests its significant role in gene expression regulation. Both QGRS Mapper and G4 Hunter score values indicate that this PQS has a remarkable potential for G4 folding with extraordinary structural stability, being higher than those of well-known G4 structures.[33] On the other hand, the analysis of the 5’-UTR region of SNCA mRNA revealed a promising sequence (mSNCA) that alone contains nearly 46% of previously mutated guanines to validate the significance of mRNA G4 structures linearization in SNCA translation inhibition.[34] ChIP and G4-RIP analyses were used to confirm respectively the folding of pSNCA and mSNCA G4 structures in differentiated SH-SY5Y cells, a well-known and commonly used neuronal cell type for neurobiology research.[56] qPCR stop assay demonstrated DNA polymerase inhibition, with target amplification decreasing in direct proportion to an increase in potassium concentrations. These findings suggest the formation of G4 structures, whose stabilization increases as potassium levels rise, that act as a geometric obstacle to polymerase.[57] Biophysical studies on model oligonucleotides compellingly demonstrate that pSNCA and mSNCA sequences fold into very stable G4 structures, highlighting their potential as therapeutic targets in neurodegenerative diseases. Specifically, pSNCA (G_4_ATG_4_CAG_5_CGCG_4_, 24nt) sequence adopted a mainly antiparallel G4 topology with high thermal stability, reaching over 95 °C at physiological potassium concentrations and around 60 °C at 1 mM K^+^. pSNCAext (GATG_4_ATG_4_CAG_5_CGCG_4_TGA, 30-nt) sequence was also tested to better understand how the neighboring bases influence pSNCA G4 folding. The addition of flanking nucleotides altered the equilibrium to an equally stable G4 with a hybrid topology. These results demonstrate that both pSNCA and pSNCAext are very stable in comparison with other highly structured DNA-G4 models, with melting temperature values around 12 degrees higher than the one identified for the most stable parallel G4 known, Myc-23456, in the presence of 5 mM KCl. Additionally, mSNCA ((G_2_A)_3_CUA(G_2_A)_3_G_2_, 23 nt) parallel folding was confirmed, indicating good RNA-G4 stability with a melting temperature of around 65°C.

The use of well-known G4-stabilizing ligands (HPHAM and PDS) emphasized the prospect of enhancing SNCA-G4s stability leading to the modulation of SNCA transcription and translation *in vitro* in the SH-SY5Y cell line. HPHAM resulted in the best-performing stabilizing ligand by biophysical assay due to its remarkable capacity to conform closely to the original structure of the pSNCA G4, without inducing distortions to increase interaction. In contrast, TMPyP4 destabilized DNA-SNCA G4s pSNCA and pSNCAext and slightly stabilized RNA-SNCA G4 mSNCA, in line with its controversial behavior in interacting with G4 structures.[47] Fascinating, SH-SY5Y cells’ treatment with HPHAM led to an increase in SNCA expression as a result of novel RNA synthesis, aligning with the recent findings that G4 formation and stabilization can actively contribute to enhancing, rather than repressing, gene expression.[21] Nevertheless, both HPHAM and PDS significantly reduced α-Syn protein production by over 50%. Considering their effect on SNCA mRNA G4, this promising protein reduction is likely due to the impairment of the translation process caused by HPHAM- or PDS-induced stabilization of mSNCA in the 5’-UTR of SNCA mRNAs.[34] This assumption is reasonable as both HPHAM and PDS are known to be excellent generic binders for G4s structures, suggesting they can effectively bind the G4 structures within SNCA mRNA inhibiting α-Syn translation. Although it has demonstrated that the presence of G4 structures in the SNCA 5’-UTR mRNA region mediates a negative response on SNCA mRNA translation,[34] it remains to be defined how the newly discovered G4 on the promoter (pSNCA) interferes with transcription. Therefore, a PNA sequence designed to hybridize the core nucleobases of pSNCA sequence (NLS-PNA-C343) was engineered to successfully drive the unfolding of the highly stable G4 within the SNCA gene’s TSS. The disruption of pSNCAext-G4 by NLS-PNA-C343 favored the formation of a stable PNA-DNA heteroduplex at the physiologic temperature of 37°C. PNAs, unlike other antisense tools, allow for a tighter bond with natural nucleic acids due to the lack of polyanionic electrostatic interactions (the backbone of PNAs is an achiral aminoethyl glycyl spacer) and greater stability as they are not recognized by nucleases or proteases.[58] Furthermore, PNAs can efficiently discriminate a single base mutation with remarkable specificity, making this tool highly selective, as demonstrated by the results obtained with the five-bases mismatched NLS-PNA5. [49] The selective targeting of the TSS G4 in the SNCA gene promoter by NLS-PNA-C343 conjugate led to a significant decrease in both mRNA and protein levels, approximately 70%. This result strongly suggests that the specific targeting and disruption of this interesting and newly discovered G4 within the SNCA promoter is enough to successfully down-regulate both SNCA mRNAs and α-Syn protein expression. The same result cannot be expected from TMPyP4. Despite its biophysical data indicating the destabilization of the target of interest, the latter lacks the necessary specific selectivity towards pSNCA G4 to effectively reduce α-Syn protein production.

To summarize, this work demonstrates that at the steady state in physiological conditions (Figure 5A) the new pSNCA DNA-G4 is involved in the normal regulation of SNCA expression. In this interplay, the G4 located in 5’UTR (mSNCA) controls a-syn translation. The stabilization of pSNCA DNA-G4 folding with highly-performance G4 ligands increased mRNA production (Figure 5B). Nevertheless, the non-specificity of these ligands suggests that they may also stabilize G4s in the SNCA 5’-UTR mRNA region, leading to translation impairment and a lowering in α-Syn protein levels, consistent with previous data (Figure 5B).[34] More attractive, the selective disruption of pSNCA DNA-G4 efficiently inhibited the SNCA transcription, with a consequent decrease in α-Syn expression resulting from fewer mRNAs (Figure 5C). Considering the high stability of pSNCA G4, along with its unique localization straddling the TSS, our results support the potential of this new strategy in restoring healthy α-Syn levels within neurons of neurodegenerative disease-affected patients. Although the development of new hyper-selective ligands for these innovative SNCA-G4s is of pivotal importance, these data, with unprecedented nucleic acid targets and their innovative mechanism of action, offer a novel mechanistic perspective for the possible future development of personalized therapeutic strategies against synucleinopathies.

**Figure 5.**
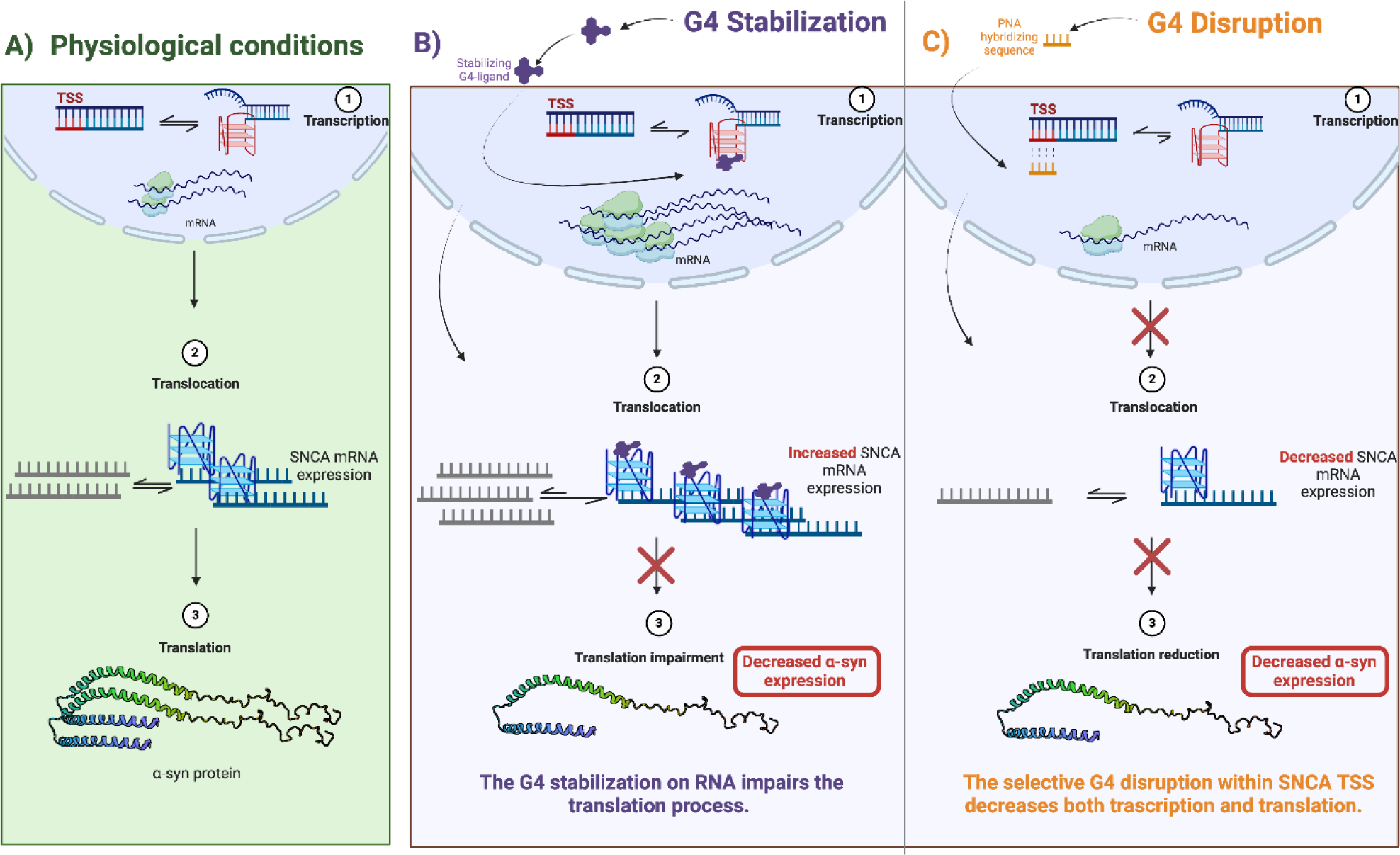
Schematic illustration of the proposed mechanism of action of SNCA G4s under investigation. Starting from physiological conditions (panel A), the stabilization of the pSNCA G4 (within SNCA TSS) causes the accumulation of SNCA mRNA, parallelly the mSNCA G4 stabilization (in the 5’-UTR mRNA of SNCA) impairs protein translation and consequently reduces α-syn expression (panel B). Instead, the specific pSNCA G4 disruption decreases the transcription process and causes the reduction of α-syn levels (panel C).

## EXPERIMENTAL SECTION

Solvents, salts, BRACO19 hydrochloride, and TMPyP4 were purchased from Merck and TCI Chemicals and were used as they were without further purification. HPHAM,[59] Pyridostatin,[60] and C343[61] were synthesized according to the previously reported protocols, and their identity was assessed by ^1^H-NMR. Analytic HPLC profiles were recorded to confirm HPHAM and Pyridostatin purity major of 98% (Figures S16 and S17). For their chemical structure, please see Figure S18.

Fmoc-protected amino acids and resin for SPPS were bought from Novabiochem® (Merck, Milan, Italy), while Fmoc/Bhoc-protected PNA monomers were from Panagene (Daejeon, South Korea).

### Oligonucleotides

All the model oligonucleotides were purchased from Eurogentec (Seraing, Belgium) and Biomers.net GmbH (Ulm, Germany) with an HPLC grade of purity (See Table S5 for sequence details).

They were dissolved in nuclease-free water and stored at -20°C. Exact oligonucleotide strand concentrations were determined by absorption measurements at 260 nm, using the extinction coefficient values provided by the manufacturer.

### PNA-conjugates synthesis

PNA-conjugates were sequentially assembled using microwave-assisted Fmoc solid phase peptide synthesis on a semi-automatic synthesizer (Biotage® Initiator+). The synthesis was conducted on a Rink Amide resin (100-200 mesh; loading 0.37 mmol/g) operating in a 0.06 mmol scale. It was pre-swelled in DCM and DMF for 1 h each and Fmoc-deprotected using 25% (v/v) piperidine in DMF. Coupling reactions for NLS synthesis were conducted at 50°C for 15 minutes, using 3 equivalents of Fmoc-protected amino acids, 3 equivalents of HOBt, 3 equivalents of PyBOP, and 6 equivalents of DIPEA in a 1:1 DMF and DCM mixture. To assemble the PNA sequence, coupling steps were carried out for 35 min at 75°C with 3 equivalents of Fmoc/Bhoc protected PNA monomers, 3.5 equivalents of Oxyma Pure®, and 3.5 equivalents of N,N′-diisopropylcarbodiimide in 1:1 NMP and DCM mixture. All the Fmoc-deprotection steps were done by treating the resin twice (3 and 10 min) with a 25% piperidine solution in DMF at room temperature.

For NLS-PNA-C343, Coumarine 343 was directly conjugated on the solid phase at room temperature using 5 equivalents of HATU and 10 equivalents of DIPEA in DMF, by stirring the mixture for 2h. Resin cleavage was performed using a TFA:TIS:MilliQ H_2_O (95:2.5:2.5, v/v) solution. Following three hours of stirring at room temperature, the resin was filtrated under low pressure, and cold diethyl ether was added to induce the precipitation of PNA-conjugates. The crudes were resuspended in acidic water (0.1% TFA) and purified by preparative HPLC (Agilent Technologies 1260 Infinity HPLC fitted with a diode array detector) using a Waters XSelectHSS C18 (2.5 µm, 50 x 4.6 mm) column. Eluents: acidic water (0.1% TFA) and acetonitrile. Purification procedure: 2 min of 95% acidic water, 14 min of gradient step to reach 60% aqueous phase; 30 mL/min flow rate; λ = 210 nm. The purity of PNA-conjugates conjugate was evaluated by analytic HPLC, resulting equal to 97.6% for NLS-PNA-C343 (Figure S19) and 98.5 % for NLS-PNA5 (Figure S20).

PNA conjugates were characterized using a surveyor UHPLC system (Thermo Finnigan, San Jose, CA, USA) equipped with a BEH Acquity UPLC column (1.7 µm) 2.1 x 50 mm, and an LCQ ADV MAX ion-trap mass spectrometer, with an ESI ion source (Figures S21-S22).

#### NLS-PNA-C343

MW: 3651.9 u.m.a.; UHPLC-MS (positive mode): 913.68 [NLS-PNA-C343+4H]^4+^; 731.22 [NLS-PNA-C343+5H]^5+^; 609.75 [NLS-PNA-C343+6H]^6+^; 522.85 [NLS-PNA-C343+7H]^7+^; 457.45 [NLS-PNA-C343+8H]^8+^ m/z + signals related to conjugates with TFA.

#### NLS-PNA5

MW: 2678.5 u.m.a.; UHPLC-MS (positive mode): 894.00 [NLS-PNA5+3H]^3+^; 670.67 [NLS-PNA5+4H]^4+^; 537.00 [NLS-PNA5+5H]^5+^; 447.58 [NLS-PNA5+6H]^6+^ m/z

### Bioinformatic prediction

SNCA promoter region (Chr4: 89835200-89838801; ENSR00000170730) and human SNCA mRNA (NM_000345.3) were analyzed by online available algorithm-based software *QGRS Mapper,*[29] *G4 Hunter,*[30] and *QuadBase2* [31] for predicting putative G4 sequences in both coding and non-coding strands. For *QGRS Mapper* analysis, were considered the following restrictions: maximum length of 30 nt; minimum G-Group size of 2 nt; loop size of 0−15 nt. For *G4 Hunter* calculations have been set at a threshold of 1.5 with a window size of 20 nt. QuadBase2 evaluation has been configured with a medium stringency (G3L1-7), considering stem length of 3 nt, loop size of 1-7 nt, and a Greedy algorithm type.

### Circular Dichroism studies

Circular Dichroism analyses were recorded by JASCO J-1500 spectropolarimeter (JASCO corporation, Easton, MD, USA), equipped with a Peltier temperature controller, in a quartz cuvette with a 10 mm optical length.

All oligonucleotides were diluted from stocks to the final concentration (2.5 µM) in lithium cacodylate buffer (10 mM, pH 7.4) containing different concentrations of KCl (0, 0.5, 1, 5, 10, 50, 100 mM). Ionic strength was kept constant by adding suitable quantities of LiCl (to 100 mM chloride salts). The solutions were annealed by heating at 95°C for 5 min and gradually cooling to room temperature over 4h.

CD spectra were recorded at 293 K, an instrument scanning speed of 200 nm/min with a response of 2 s over a wavelength range of 210-340 nm with a 1 nm sampling interval. The reported spectra of each sample represent the average of 4 scans and are baseline-corrected for signal contributions due to the buffer mixture. CD spectra were normalized to Δε (M^-1^cm^-1^) according to the relationship Δε = ϑ / (32980 x C x l), where ϑ corresponded to observed ellipticities (in millidegrees), C is DNA molar concentration, and l is the path length in centimeters. Secondary and tertiary structure fractions predictions were calculated by exploiting the algorithm developed by Del Villar-Guerra et al.[36] For the CD-melting experiments of G4 structures (2.5 µM), the ellipticity was recorded at 295 nm for pSNCA and pSNCAext and at 264 nm for mSNCA every 0.5°C, with a temperature scan rate of 1 °C/min in the range of 15-95 °C, after a pre-equilibrium of the solution at 15°C for 5 minutes. Data were the average of three independent experiments. G4-ligands were added in 10 µM concentration (4 equivalents) and left to equilibrate for 16 h at room temperature before recording the melting curve. PNA conjugates were added in 10 µM concentration (4 equivalents) and left to equilibrate for 4 h at room temperature before recording the CD spectrum and the melting curve. All the melting curves are reported in the Supporting Information. Melting temperature values were calculated according to the Van’t Hoff equation, applied for a two-state transition from a folded to unfolded state, assuming that the heat capacity of the folded and unfolded states are equal. ΔT_m_ was calculated as the difference between T_m_ in the presence and absence of the compounds.

### UV-vis studies

UV-visible spectra were run on an Agilent Cary 60 spectrophotometer (Agilent Technologies, Santa Clara, CA, USA) equipped with a Single-Cell Peltier temperature controller using a quartz cuvette of 10 mm path length. pSNCA and pSNCAext model oligonucleotides were diluted from stocks to final concentration (2.5 µM) in lithium cacodylate buffer (10 mM, pH 7.4) with 1 mM or 5 mM KCl. Ionic strength was kept constant by adding suitable quantities of LiCl (to 100 mM chloride salts). mSNCA was suspended in the same conditions but in the presence of 10 mM or 100 mM KCl. The solutions were annealed by heating at 95°C for 5 min and gradually cooling to room temperature over 4h. The absorbance was recorded as a function of temperature (20-95°C) from 220 to 340 nm wavelength intervals at a scan rate of 50 nm/min and a slit width of 0.5 nm. All the melting curves are reported in the Supporting Information. Mid-transition temperatures were determined following the absorbance decreasing at 295 nm in the function of temperature (20-95°C). Data were the average of three independent experiments. Model-dependent thermodynamic parameters were estimated by following the protocol published by J.L. Mergny and collaborators [42]. In particular, the association constants (Ka) were calculated in function of the folded fractions (ø_T_), accordingly the equation Ka = ø_T_/(1-ø_T_). Van’t Hoff enthalpy and entropy values were estimated accordingly the equation LogKa = (−ΔH°/R) x ((1/T) + ΔS°/R), by restricting the temperature range for which folded fraction is 0.15 <ø < 0.85. R, the perfect gas constant, was considered equal to 1.987 calmol^-1^K^-1^. UV-vis thermal difference spectra have been determined by the mathematical subtraction of the spectra of fully unfolded oligonucleotides (above Tm, 95°C) and the spectra of fully folded oligonucleotides (low T, 20°C) [42]. TDS factors correspond to the average of ΔA_240nm_/ΔA_295nm_ absolute values obtained at 20, 30, 40, and 50 °C.

### FRET-melting

FRET-melting measurements were recorded by an Agilent AriaMx Real-time PCR System (Agilent Technologies, Santa Clara, CA, USA) using 5’-FAM (6-carboxyfluorescein) and 3’-TAMRA (6-carboxy-tetramethylrhodamine) labeled oligonucleotides.[62] In a total volume of 60 µL, 0.25 µM of tagged oligonucleotides were dissolved in 10 mM lithium cacodylate buffer pH 7.4 in the presence of proper KCl/LiCl concentrations (suitable quantities of LiCl were added to maintain the total concentration of chloride salts equal to 100 mM). The mixtures were annealed by heating at 95 °C for 3 min followed by an equilibration step in ice for 30 min. The folded samples were stored at -4°C. Melting experiments were performed on 96 wells plates, by analyzing 20 µL of samples. After the first equilibration step at 25 °C for 5 minutes, a stepwise increase of 1°C/min was performed to reach 95°C, measuring the fluorescence emission at 516 nm (excitation at 462 nm) according to SYBR/FAM optical cartridge (Agilent). Data were the average of three independent experiments, each conducted in triplicate conditions. G4-ligands were added in 1 µM concentration (4 equivalents) and left to equilibrate for 16 h at room temperature before recording the FRET-melting curve. The fluorescence emission curves were normalized between 0 and 1, and T_m_ was defined as the temperature corresponding to 0.5 normalized emissions. ΔT_m_ was calculated as the difference between T_m_ in the presence and absence of the compounds. The melting curves are reported in the Supporting Information.

### NMR studies

^1^H-NMR spectra were recorded on Bruker Avance NEO 700MHz (BRUKER, Billerica, MA, USA), equipped with a triple resonance helium-cooled cryoprobe 5 mm. In a total volume of 800 µL, 0.17 mM of oligonucleotides were dissolved in 90% H_2_O/10% D_2_O solution with 100 mM KCl and 20 mM potassium phosphate buffer at pH 7.0. The mixtures were annealed by heating at 95 °C for 5 min followed by an equilibration step in ice for 30 min. The folded samples were stored at -4°C. ^1^H-NMR spectra were recorded at 25°C using a zgesgp technique to suppress the water signal. All NMR spectra were processed and analyzed using Bruker TopSpin 2.1.

### Chemometric analysis on CD spectra

Principal Component Analysis (PCA) was performed using R software in which the Chemometric Agile Tool was loaded. PCA was initially applied to a reference dataset of 23 G4s CD spectra (7 parallels, 8 hybrids, and 8 antiparallels) reported in the literature,[36] by considering only the first two components PC1 and PC2. The variables were centered and scaled. Calculated scores from pSNCA, pSNCAext, and mSNCA spectra (recorded at 2.5 µM oligonucleotides, 100 mM KCl, and 10 mM lithium cacodylate buffer, pH 7.4) were projected on the training set. The dataset was then utilized to examine the impact of NLS-PNA-C343 and NLS-PNA5 on the pSNCAext structure (spectra recorded after the addition of 4 equivalents of PNA conjugates and an equilibration time of 4h).

### SH-SY5Y culture and maintenance

SH-SY5Y cells were purchased from ATCC. Cells were maintained in DMEM/F12 Complete Medium supplemented with 10% of Fetal Bovine Serum (FBS; Microtech), 1% glutamine (L-GLUT; GIBCO), and 1% Penicillin/Streptomycin (P/S, GIBCO) at 37 °C in a 5% CO_2_. For all performed experiments, cells were differentiated with retinoic acid (RA) at a concentration of 10 μM for 7 days in a 1% FBS DMEM High Glucose medium to induce differentiation.

### Chromatin Immunoprecipitation (ChIP)

ChIP G4-immunoprecipitation was performed on SH-SY5Y cells following a previously published protocol.[16] Cells were fixed in 1% formaldehyde for 10 minutes and quenched with 0.125 M glycine. Chromatin was sonicated using Bioruptor® plus sonicator (Diagenode) to shear to an average size of 1,000–1,500 bp and incubated in the presence of BG4 (Merck, Milan, Italy, MABE817, Lot: 3534080). Anti-FLAG antibody (Sigma-Aldrich, Milan, Italy, cat. F1804) coated Protein-G magnetic beads (Pierce™ ThermoFisher Scientific) were used to capture BG4-G4 complexes. After de-crosslinking, samples were purified with the ExpIn Combo GP mini kit (GeneAll). G4 enrichment was quantified via qPCR, using the Optimum QMastermix with SYBER GREEN (Genespin) and with the CFX Connect Real-Time PCR System (Bio-Rad). GAPDH was used as a negative control, primer sequences are reported in Table S6. Relative enrichments were calculated with respect to their inputs.

### RNA Immunoprecipitation

RNA G4-immunoprecipitation was performed on SH-SY5Y cells. Cells were lysed with lysis buffer (10 mM Tris HCl, 10 mM NaCl, 2 mM EDTA, 0.5 Triton X100) and incubated on ice for 10 minutes. 120mM NaCl was then added and cells were incubated for a further 5 minutes. Cells were then pelleted at 16000 g for 15 minutes and the surnated was retained for pre-clearing. Following pre-clearing, the lysate was incubated in the presence of BG4 (Merck, Milan, Italy, MABE817, Lot: 3534080). Anti-FLAG antibody (Sigma-Aldrich, Milan, Italy, cat. F1804) coated Protein-G magnetic beads (Pierce™ ThermoFisher Scientific) were used to capture BG4-G4 complexes. After elution, samples were purified with TRIZOL® reagent (Thermo Fisher Scientific) following the manufacturer’s instructions. Samples were retrotranscribed and analyzed via Real-Time PCR (see below). GAPDH was used as a negative control, primer sequences are reported in Table S6. Relative enrichments were calculated with respect to their inputs.

### qPCR-Stop assay

qPCR stop assay was used to determine whether the putative genomic pSNCA region folds into a G4 structure.[63] Primers were designed to surround the G4 and increasing KCl concentrations (0, 0.1 mM, 1 mM, 10 mM, and 50 mM) were added to stabilize the G4 and thus assess Taq activity. Real-Time PCR was performed with the CFX Connect Real-Time PCR System (Bio-Rad) using the Optimum QMastermix with SYBER GREEN (Genespin). Primer sequences are reported in Table S6.

### MTT assays

SH-SY5Y cells were seeded in a 96-well plate at 2×10^5^ cells/well density and they were treated with increasing concentrations of G4 ligands. 10 μL MTT assay kit reagent (Merck) was then added to each well, and cells were incubated for 3 h. MTT crystals were eluted with 100 μL of elution solution, composed of 4 mM HCl, 0.1% (v/v) NP40 all in isopropanol for 30 min. The relative absorbance was measured with a Multiskan GO spectrophotometer (Thermo Fisher Scientific) at λ = 570 nm.

### Immunocytochemistry

SH-SY5Y cells were seeded on ethanol-washed glass coverslips and cultured in a standard medium. Following treatment with HPHAM, cells were fixed with 4% paraformaldehyde in 0.1 M PBS (Thermo Fisher Scientific), pH 7.4, for 20 min at room temperature, and then washed with PBS. Nuclei were stained with DAPI at the final concentration of 1 μg/ml for 30 min. Glass coverslips were mounted using the FluorSave Reagent (Calbiochem) and analyzed by confocal microscopy (Confocal laser scanning microscopy platform Leica TCS SP8, Leica Microsystems). Co-localization rate was analyzed with Leica Analysis Software.

### RNA extraction and real-time PCR

Total RNA was extracted using TRIZOL® reagent (Thermo Fisher Scientific) following the manufacturer’s instructions and then quantified using the Multiskan GO spectrophotometer (Thermo Fisher Scientific). For stability assays, cells were co-treated with 1.5 µg/ml Actinomycin D. 1500 ng of RNA were retro-transcribed using the Super Mix 5X Full (Genespin) kit following the manufacturer’s instructions. Real-Time PCR was performed with the CFX Connect Real-Time PCR System (Bio-Rad) using the Optimum QMastermix with SYBER GREEN (Genespin). The NCBI’s Primer-BLAST tool was used to design primers. Primer sequences are reported in Table S6. Gene expression was calculated using the 2^−ΔΔCt^ method. TBP was used as endogenous control.

### Western blotting analysis

Protein extracts were obtained by means of RadioImmunoPrecipitation Assay (RIPA) lysis buffer, proteins were quantified with the Bradford Assay following a standard protocol (Coomassie Plus— The Better Bradford Assay™ Reagent, Thermo Scientific), and equal amounts of solubilized proteins were heated in Laemmli sample buffer (Bio-Rad) containing 70 mM 2-β-mercaptoethanol (Sigma Aldrich). Samples were separated in a 12% SDS-PAGE gel and electroblotted onto a nitrocellulose membrane (Trans-blot, Bio-Rad). Membranes were then fixed with 4% paraformaldehyde and 0.01% Glutaraldehyde in PBS (Thermo Fisher Scientific), for 30 min at room temperature, and then blocked in 5% slim milk (diluted in TBS with 0.05% Tween-20). After that, membranes were probed with the appropriate primary antibody: Recombinant Anti-Alpha-synuclein antibody [MJFR1] (1:2000 Abcam, ab138501) or β-Actin (1:5000 Sigma-Aldrich, A1978), overnight at 4°C. Lastly, membranes were incubated with the appropriate secondary antibody Peroxidase AffiniPure Goat Anti-Rabbit/Mouse IgG (1:5000 dilution; Jackson Immuno Research). Proteins were visualized by means of an enhanced chemiluminescence detection solution (Thermo Fisher Scientific). After acquisition by a GelDoc™ image capture system (Uvitec, Eppendorf), densitometric analysis of the bands was performed using the ImageJ software.

## Supporting information

Supporting Information_SNCA

## SUPPLEMENTARY INFORMATION

The online version contains supplementary material. %GC Content distribution; putative G-quadruplex sequences identified; estimation of the fractions of tertiary and secondary structural elements; melting data; thermodynamic parameters; qPCR stop assay for the negative control; CD spectra of pSNCAext/NLS-PNA-C343 hetero duplex and for pSNCAext/NLS-PNA5 conjugates; PCA score plot; MTT assay; immunofluorescence assay; real-time PCR of different housekeeping genes; analytic HPLC profiles; chemical structures of HPHAM, PDS, BRACO19, TMPyP4, NLS-PNA-C343, and NLS-PNA5; name and nucleobases sequences of the DNA model oligonucleotides analyzed; LC-MS analysis of NLS-PNA-C343 and NLS-PNA5; List of primers used in this study; Melting profiles.

## AVAILABILITY OF DATA AND MATERIALS

The datasets supporting the conclusions of this article are included within the article (and its additional file). Additional material or information can be provided upon request by contacting the corresponding author.

## Author Contributions

Conceptualization, Methodology and Supervision: V.P., S.C.; Formal analysis, Investigation: V.P., F.R., V.F., L.E., E.M., C.P., R.D.G.; Validation, Data Curation: V.P., F.R., V.F., M.M., O.P., P.G., M.F., S.C., C.C.; Writing – Original Draft: V.P., F.R.; Writing – Review & Editing: V.P., F.R., P.G., M.F., S.C.; Funding acquisition: V.P., F.D., M.F., S.C.; C.C.. All authors have given approval to the final version of the manuscript.

## Funding Sources

This work was supported by a Postdoctoral research fellowship (V.P.) [DCHIMICA2020-A01] from the University of Pavia, Pavia, Italy.

## ABBREVIATIONS

α-Syn: Alpha-synuclein
PD: Parkinson’s disease
G4(s): G-quadruplex(es)
TSS: transcription start sites
UTR: untranslated region
NLS: nuclear localization sequence
PNA: peptide nucleic acid
TDS: thermal difference spectra
Tm: mid-transition temperature
C343: Coumarin 343
PCA: Principal Component Analysis.

## Notes

### Competing Interest Statement

The authors have declared no competing interest.

### Summary of Updates

In-cell experiments with TMPyP4 were added to verify the contribution of a G4-destabilizing ligand (considering the biophysical data on pSNCA). Firstly, the treatment of differentiated SH-SY5Y cells with TMPyP4 did not significantly affect SNCA mRNA levels, as verified by Real-Time PCR (Figure S11A). In contrast, it drastically decreased MYC mRNA levels (Figure S11B). Additionally, TMPyP4 did not significantly affect either α-Syn expression levels if not after 48 hours, which however is a constant behavior for all the binders tested (Figure S13). The authors suggest that this behavior could be ascribed to the non-specificity of the ligand, which interacts with both pSNCA and mSNCA G4s. Its destabilizing properties on pSNCA are insufficient to reduce SNCA mRNA production, as well as its stabilizing effect on mSNCA G4 (please refer to Table 2 in the main text) did not lead to a significant decrease in α-Syn production over 24 hours. Additionally, to verify the specific contribution of the PNA structure in lowering the production of α-Syn, the authors added experiments with a sequence with five base mismatches with pSNCAext to emphasize the specificity of the tool. CD spectrum analysis revealed that NLS-PNA5 did not induce a significant change in the pSNCAext structure, mainly maintaining the G4 hybrid topology (Figure S6). This behavior was also confirmed by PCA analysis (Figure S4). Furthermore, treatment of differentiated SH-SY5Y cells with non-toxic NLS-PNA5 (the same concentration used for NLS-PNA-C343 for comparison, see MTT assay in Figure S15A) did not affect either mRNA levels or α-Syn production (Figure S15 B and C). These results emphasized that PNAs can efficiently discriminate a single base mutation with remarkable specificity, making these tools highly selective.

