## Supporting Information_SNCA for "Effective lowering of α-synuclein expression by targeting G-quadruplex structures within the SNCA gene": 2024-07-25_Pirota et al_SNCA_supp Info_revised.pdf

‡ Joint authors.

**TABLES OF CONTENTS** Pages


| | |
|---|---|
| **Figure S1.** %GC Content distribution in the SNCA promoter region | S1 |
| **Table S1.** Putative G-quadruplex sequences identified in the SNCA promoter region | S1 |
| **Figure S2.** qPCR stop assay on SH-SY5Y genomic DNA for the negative control | S2 |
| **Figure S3.** Estimation of the fractions of tertiary and secondary structural elements of SNCA G4s | S2 |
| **Table S2.** Comparison of melting data obtained by CD-melting experiments | S3 |
| **Table S3.** Model-dependent thermodynamic parameters per SNCA G4s | S4 |
| **Table S4.** Melting temperature values resume | S4 |
| **Figure S4** PCA scores plot | S5 |
| **Figure S5.** Estimation of the tertiary structural elements‘ fractions induced by G4-ligands | S6 |
| **Figure S6.** CD spectra of pSNCAext/NLS-PNA-C343 hetero duplex | S6 |
| **Figure S7.** MTT assay following HPHAM treatment | S7 |
| **Figure S8.** MTT assay following PDS treatment | S7 |
| **Figure S9**. MTT assay following TMPyP4 treatment | S8 |
| **Figure S10.** Immunofluorescence assay shows HPHAM entrance in the cells | S9 |
| **Figure S11.** Real-time PCR after TMPyP4 treatment | S9 |
| **Figure S12.** Real-time PCR of different housekeeping genes | S10 |
| **Figure S13.** Western Blot for α-Syn quantification after TMPyP4 treatment | S10 |
| **Figure S14**. MTT assay following NLS-PNA-C343 treatment | S11 |
| **Figure S15.** Treatment of SH-SY5Y cells with NLS-PNA5 | S11 |
| **Figure S16.** Analytic HPLC profile of HPHAM | S12 |
| **Figure S17.** Analytic HPLC profile of PDS | S12 |
| **Figure S18.** Chemical structures of HPHAM, PDS, BRACO19, and TMPyP4 | S13 |
| **Table S5.** Name and nucleobase sequences of the DNA model oligonucleotides | S13 |
| **Figure S19.** Analytic HPLC profile of NLS-PNA-C343 | S14 |
| **Figure S20.** Analytic HPLC profile of NLS-PNA5 | S14 |
| **Figure S21.** LC-MS analysis of NLS-PNA-C343 | S14 |

**Figure S22.** UHPLC-MS data of NLS-PNA5 S15

**Table S6.** List of primers used in this study S16

**Denaturation curves obtained by CD-melting experiments** S17-S21

**Denaturation curves obtained by UV-melting experiments** S22

**Denaturation curves obtained by FRET-melting experiments** S23-S25

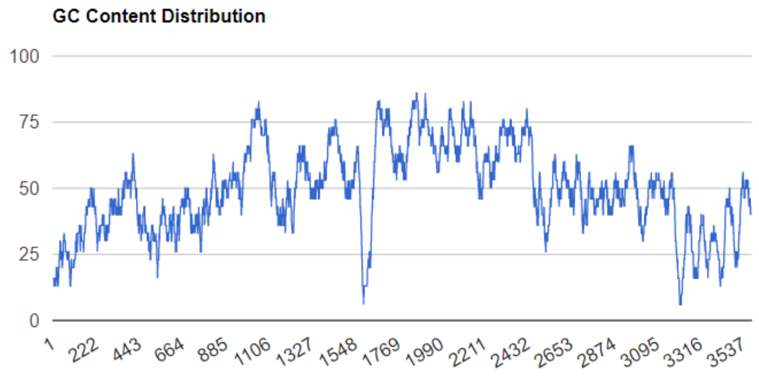


**Figure S1.** %GC Content distribution in the SNCA promoter region. Position 1 corresponds to nucleobase 89835200 on Chr4. Full Length(3602bp) | A(25.7% 926) | T(24.9% 897) | G(22.4% 808) | C(27.0% 971).

**Table S1.** Putative G-quadruplex sequences identified in the SNCA promoter region (Chr4: 89835200-89838801) with their G-Score values. (*position 1 corresponds to nucleobase 89835200 on Chr4). pSNCA sequence is highlighted in violet.

| **Position*** | **Length** | **Sequence** | **QGRS Mapper G-score** | **G4Hunter score** |
|---|---|---|---|---|
| 825 | 26 | GGAAGCTGGAGGCTTAGGGTCAGGGG | 19 | 1.35 |
| 877 | 25 | GGCTAACAGGTTGATGGTGGAAAGG | 20 | 0.76 |
| 932 | 30 | GGTGTGGTTCAAACTCAGGCAAACAGCAGG | 14 | 0.40 |
| 1289 | 19 | GGGGAAAAGCGGATCCCGG | 15 | 0.789 |
| 1638 | 29 | GGAAAGAAGAAGGAAAAGGAGCGCACAGG | 16 | 0.586 |
| 1670 | 11 | GGCGGAGGCGG | 21 | 1.27 |
| 1688 | 22 | GGGGGACTGTCCCCAGAGGAGG | 10 | 0.591 |
| 1736 | 14 | GGGGGAGTGGGAGG | 19 | 2.43 |
| 1802 | 25 | GGTGGAAAGGCAGAAGGCTTGAAGG | 20 | 0.8 |
| 1858 | 17 | GGAGGGGCCGGGCCGGG | 21 | 1.76 |
| **2102** | **24** | **GGGGATGGGGCAGGGGGCGCGGGG** | **62** | **2.75** |
| 2163 | 16 | GGCGGGAAAGTGGGGG | 17 | 2.06 |
| 2232 | 17 | GGAGCGGTTGGGCTAGG | 21 | 1.18 |
| 2403 | 30 | GGAGCTGCTGGAGGAGACAGGCAGCGCCGG | 20 | 0.533 |
| 2494 | 24 | GGTCACAGGTTACAACGTTAGGGG | 10 | 0.875 |
| 2574 | 24 | GGAAAGGGTCCTGAGGGTGAAAGG | 19 | 1 |
| 2621 | 21 | GGTTGAGGAGGCAGGAGAAGG | 20 | 1 |
| 3172 | 14 | GGAAGAGGAGGGGG | 18 | 2.07 |
| 3532 | 27 | GGACCAGAGCAAGAAGGTTGGACTTGG | 10 | 0.481 |

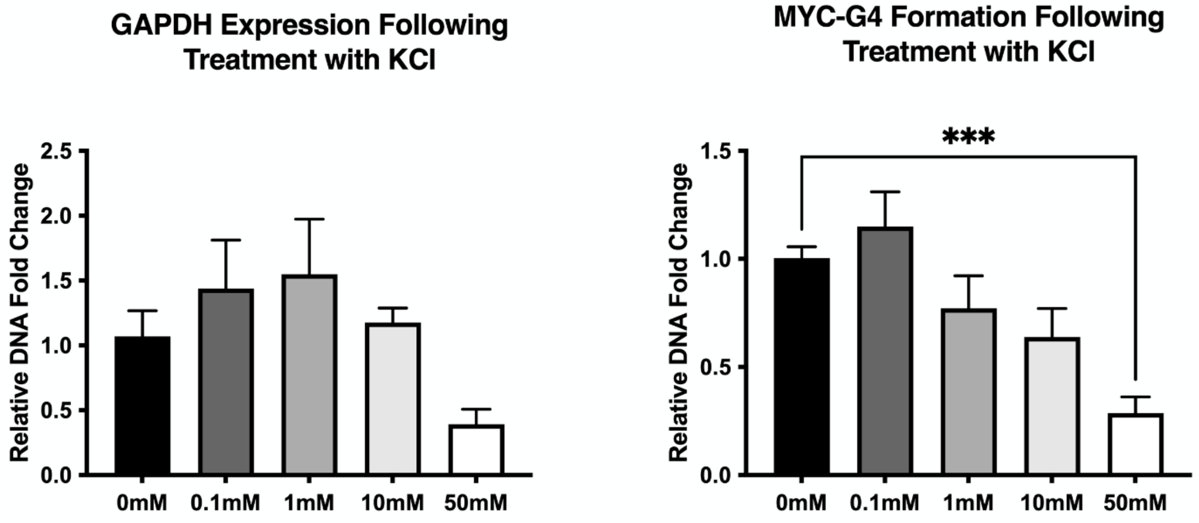


**Figure S2.** qPCR stop assay performed on genomic DNA from SH-SY5Y cells for a non-G4 forming sequence of the GAPDH gene used as negative control and for a G4 forming sequence of the MYC gene as positive control. qPCR stop assay was performed on genomic DNA obtained from SH-SY5Y after 7 days of neural differentiation. The expression of non-G4 forming GAPDH region abundance was assayed via qPCR. Data are expressed as mean ± SEM of 5 independent samples (n = 5, ***p<0.001 vs 0mM).

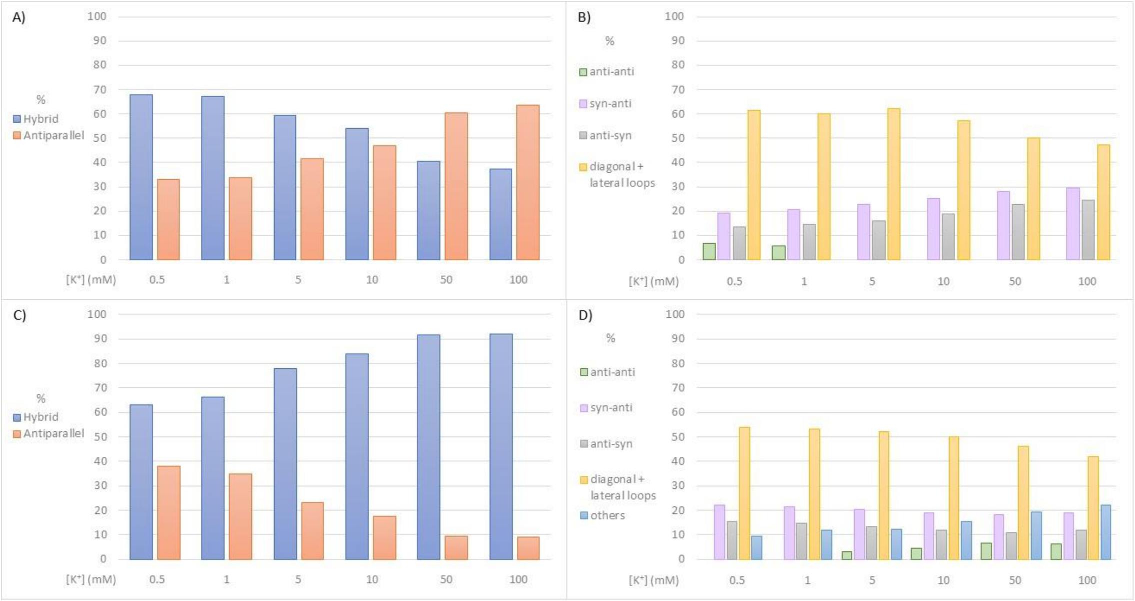


**Figure S3** – Bar graphs showing the estimation of the fractions of the tertiary structural elements of (A) pSNCA, and (C) pSNCAext G-quadruplexes, and of the secondary structural elements of (B)

pSNCA, and (D) pSNCAext G-quadruplexes. These results were derived from the CD spectra fitting by the algorithm developed by R. Del-Villar Guerra et al. [1].

**Table S2.** Comparison of melting data obtained by CD-melting experiments. KCl concentration was lowered concerning the stability of G4, to obtain a Tm value between 50-60 °C. Literature data for TERRA,[2] ADAM10,[3] and MMP16 [4] were reported.

| | **Name** | **KCl (mM)** | **Tm (°C)** |
|---|---|---|---|
| **DNA G4s** | pSNCA | 1 | 61.7 ± 0.1 |
| | pSNCAext | 1 | 56.8 ± 0.1 |
| | pSNCA | 5 | 74.6 ± 0.1 |
| | pSNCAext | 5 | 73.2 ± 0.6 |
| | Myc-23456 | 5 | 61.2 ± 0.4 |
| | cMyc | 5 | 56.3 ± 0.3 |
| | 22ag | 20 | 51.7 ± 0.6 |
| | LTR-III | 20 | 56.7 ± 0.6 |
| | CKIT-1 | 50 | 52.5 ± 0.2 |
| **RNA G4s** | mSNCA | 100 | 62.4 ± 0.1 |
| | mSNCA | 10 | 53.6 ± 0.1 |
| | TERRA [2] | 1 | 63.7 ± 0.2 |
| | ADAM10 [3] | 1 | 60 ± 1 |
| | MMP16 [4] | 10 | 69.2 ± 0.2 |

**Table S3.** Model-dependent thermodynamic parameters estimated from the thermal unfolding of pSNCA, pSNCAexr, and mSNCA model oligonucleotides by UV-melting experiments. Data processing was performed by following the protocol developed by J.L. Mergny et al. [5]. The temperature range considered was restricted for which the folded fraction is 0.15 <ø < 0.85.

| **pSNCA** | | | | | |
|---|---|---|---|---|---|
| **Values** | **KCl (mM)** | **Tm (°C)** | **ΔH° ($kcalmol^{-1}$)** | **ΔS° ($calmol^{-1}K^{-1}$)** | **$\Delta G°_{310K}$ ($kcalmol^{-1}$)** |
| UV-melting (ΔAbs 295nm) | 1 | 61.5 ± 0.4 | -55.5 ± 0.6 | -166 ± 2 | -4.02 |
| | 5 | 74.8 ± 0.3 | -86.5 ± 0.4 | -249 ± 4 | -9.31 |
| **pSNCAext** | | | | | |
| **Values** | **KCl (mM)** | **Tm (°C)** | **ΔH° ($kcalmol^{-1}$)** | **ΔS° ($calmol^{-1}K^{-1}$)** | **$\Delta G°_{310K}$ ($kcalmol^{-1}$)** |
| UV-melting (ΔAbs 295nm) | 1 | 55.1 ± 0.1 | -52.3 ± 0.9 | -159 ± 3 | -2.99 |
| | 5 | 72.8 ± 0.6 | -74.5 ± 0.3 | -217 ± 2 | -7.23 |
| **mSNCA** | | | | | |
| **Values** | **KCl (mM)** | **Tm (°C)** | **ΔH° ($kcalmol^{-1}$)** | **ΔS° ($calmol^{-1}K^{-1}$)** | **$\Delta G°_{310K}$ ($kcalmol^{-1}$)** |
| UV-melting (ΔAbs 295nm) | 10 | 53.7 ± 0.2 | -27.7 ± 0.2 | -83.8 ± 0.5 | -1.68 |
| | 100 | 62.45 ± 0.05 | -62.5 ± 0.2 | -183.6 ± 0.4 | -5.58 |

**Table S4.** Melting temperature values obtained by UV-, CD-, and FRET-melting of pSNCA, pSNCAext, and mSNCA at different KCl concentrations, in 10mM lithium cacodylate buffer pH 7.4.

| | | **pSNCA** | **pSNCAext** | | **mSNCA** |
|---|---|---|---|---|---|
| **Values** | **KCl (mM)** | **Tm (°C)** | **Tm (°C)** | **KCl (mM)** | **Tm (°C)** |
| UV-melting (ΔAbs 295nm) | 1 | 61.5 ± 0.4 | 55.1 ± 0.1 | 10 | 54.7 ± 0.2 |
| | 5 | 74.8 ± 0.3 | 72.8 ± 0.6 | 100 | 62.45 ± 0.05 |
| CD-melting (Δε 295 nm) | 1 | 61.7 ± 0.1 | 56.8 ± 0.1 | 10 | 53.6 ± 0.1 |
| | 5 | 74.6 ± 0.1 | 73.2 ± 0.6 | 100 | 62.4 ± 0.1 |
| FRET-melting (ΔF 516 nm) | 1 | 64.3 ± 0.4 | 58.8 ± 0.8 | 10 | 55 ± 1 |
| | 5 | 76.3 ± 0.6 | 74.5 ± 0.5 | 100 | 68.5 ± 0.6 |

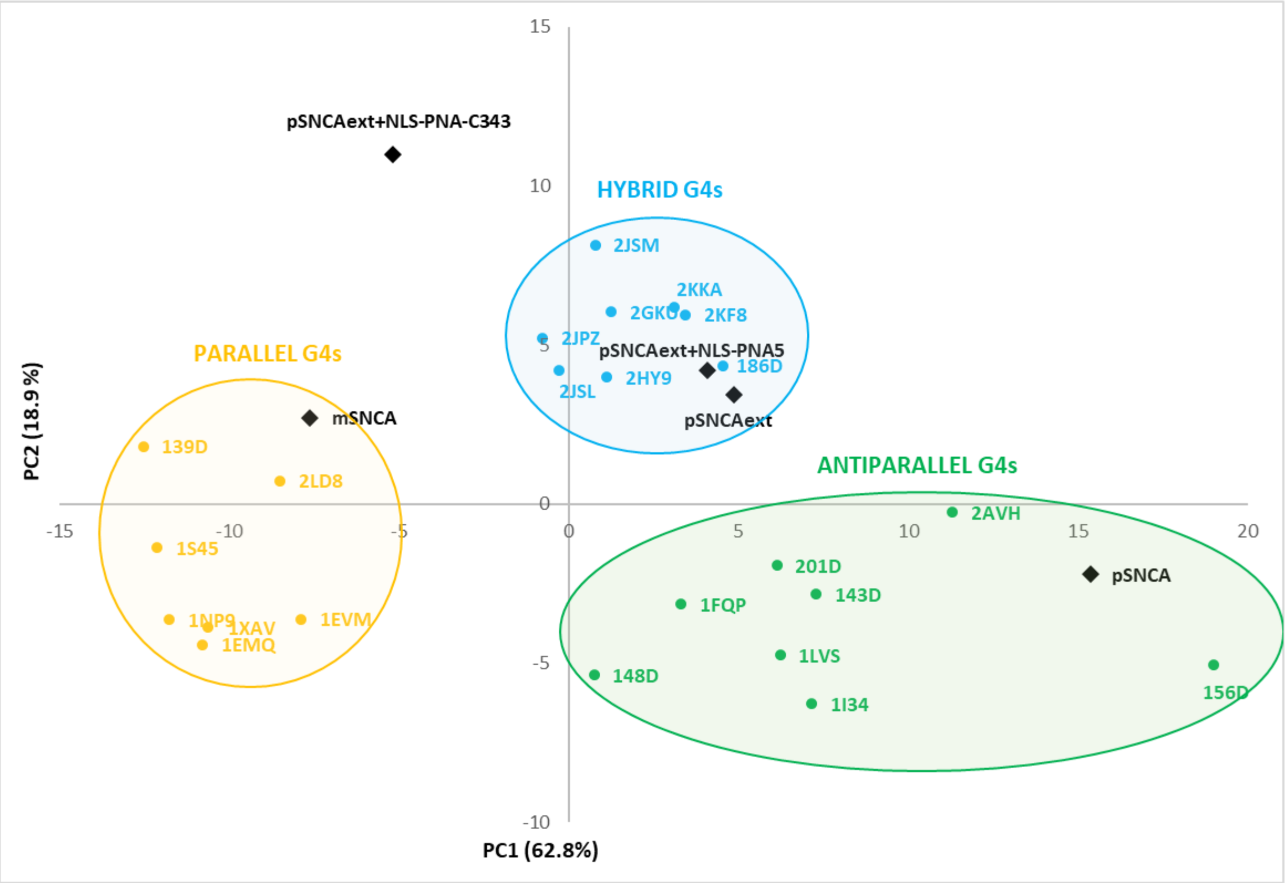


**Figure S4.** Projection of the PC1 and PC2 scores identified by the analysis of pSNCA, pSNCAext, mSNCA, pSNCAext+NLS-PNA-C343, pSNCAext+NLS-PNA5 CD spectra into the PCA score plot built on reference G4s CD spectra [1]. pSNCA, pSNCAext, and mSNCA CD spectra were recorded with 2.5 µM oligonucleotide, 100 mM KCl, and 10 mM lithium cacodylate buffer, pH 7,4, at 25°C. To the pSNCAext solution were added 4 equivalents of NLS-PNA-C343 or NLS-PNA5 (10 µM), left to equilibrate for 4h before recording the CD spectra of pSNCAext+NLS-PNA-C343 or pSNCAext+NLS-PNA5.

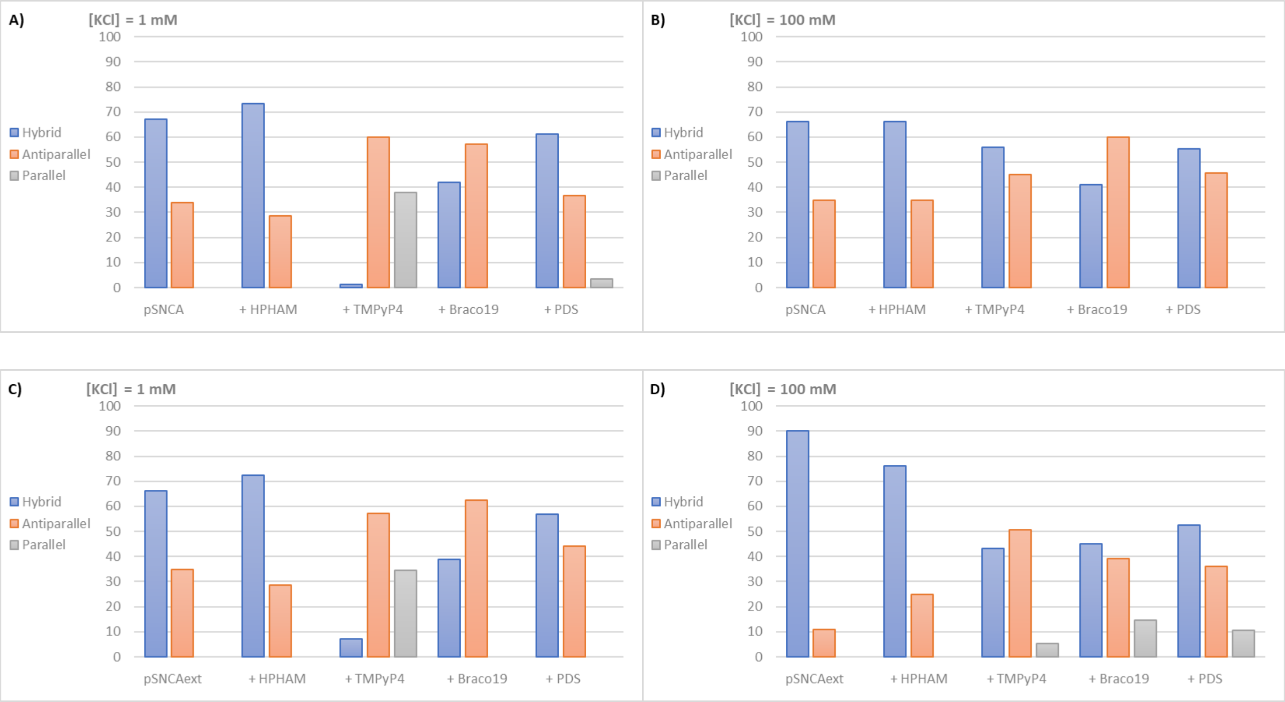


**Figure S5**. Bar graphs showing the estimation of the fractions of the tertiary structural elements of (A-B) **pSNCA** and (C-D) **pSNCAext** G-quadruplexes alone, and in the presence of 4 equivalents of well-known G4 ligands, recorded in 10 mM lithium cacodylate buffer (pH =7.4) with 1 mM or 100 mM KCl. These results were derived from the CD spectra fitting by the algorithm developed by R. Del-Villar Guerra et al. [1].

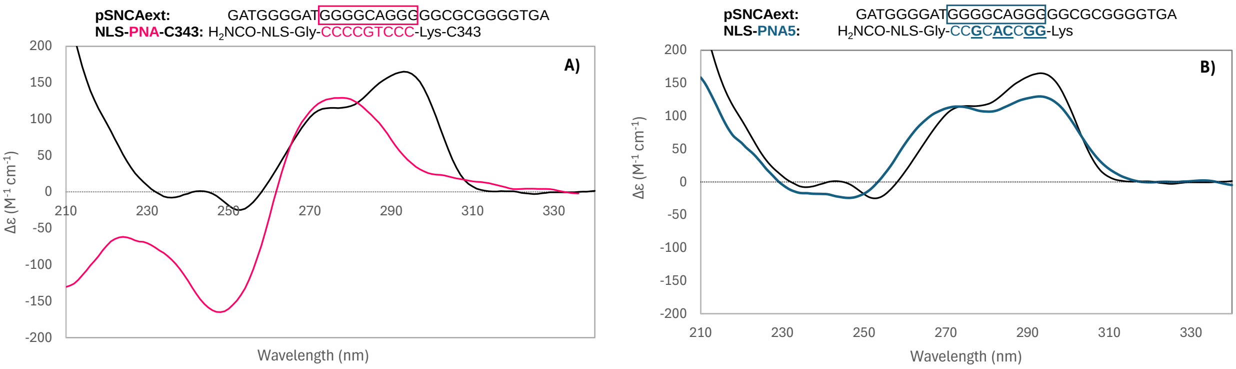


**Figure S6.** CD spectra of 2.5 µM pSNCAext alone (bold line) and in the presence of A) 10 µM NLS-PNA-C343 (pink line) or B) 10 µM NLS-PNA5 (blue line) in 100 mM KCl and 10 mM lithium cacodylate buffer pH 7.4, 25°C.

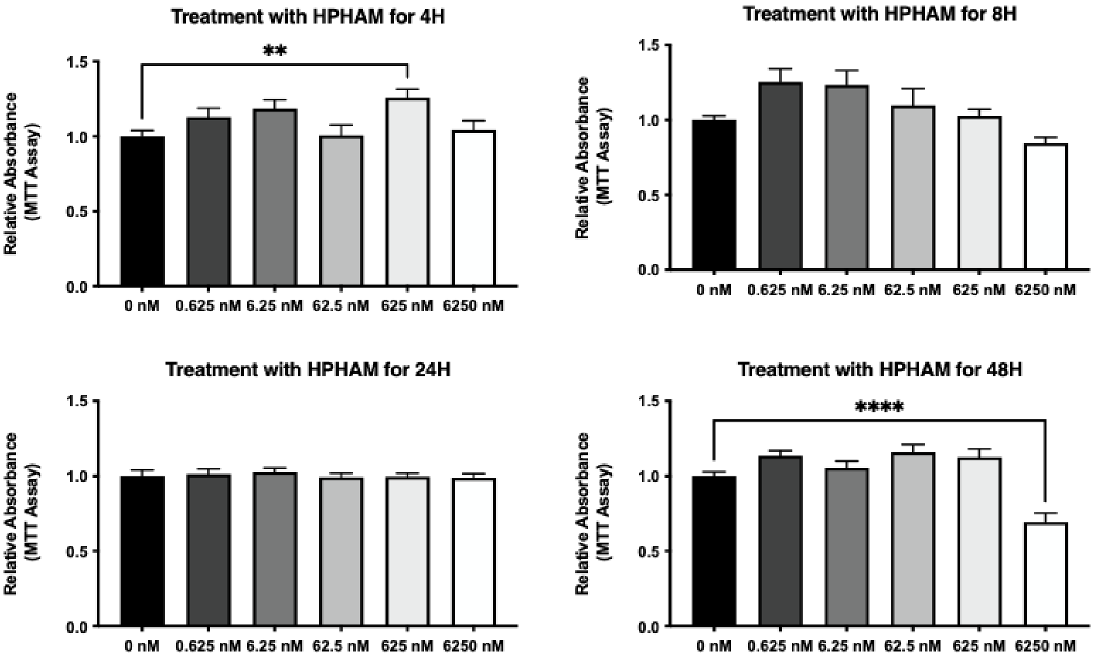


**Figure S7**. MTT assay shows the impact of HPHAM treatment on cell viability after 4, 8, 24, and 48 hours at different dosages (as reported on the x-axis). Data are expressed as mean ± SEM of 8 replicate values in 3 independent experiments (n = 24, **p<0.01, ****p<0.0001 vs 0nM)

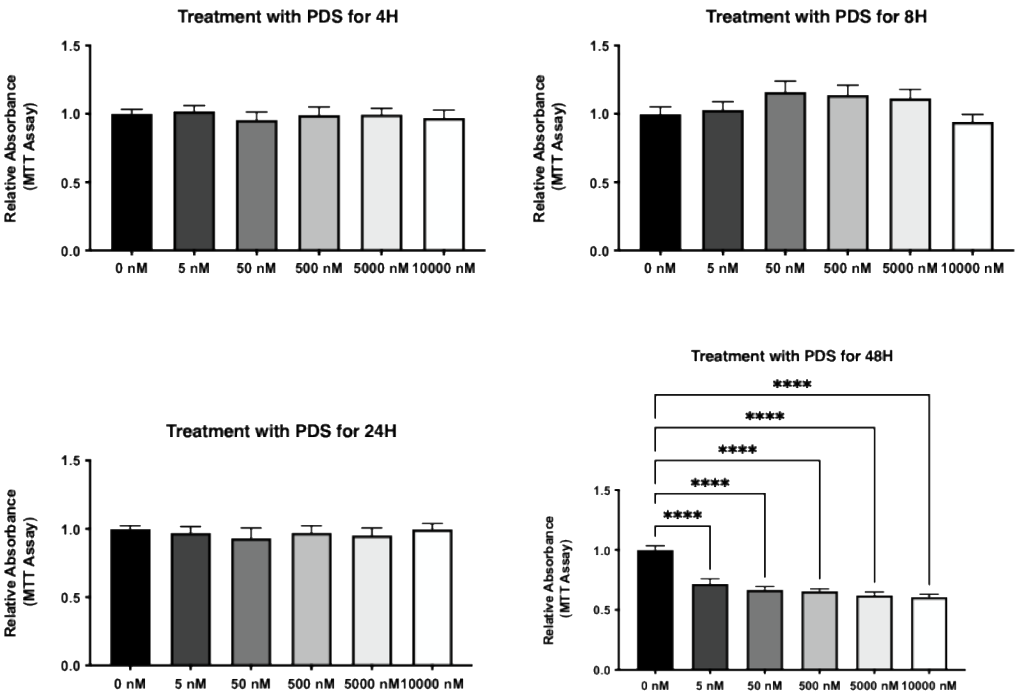

**Figure S8**. MTT assay shows the impact of PDS treatment on cell viability after 4, 8, 24, and 48 hours at different dosages (as reported on the x-axis). Data are expressed as mean ± SEM of 8 replicate values in 3 independent experiments (n = 24, ***p<0.001, ****p<0.0001 vs 0nM).

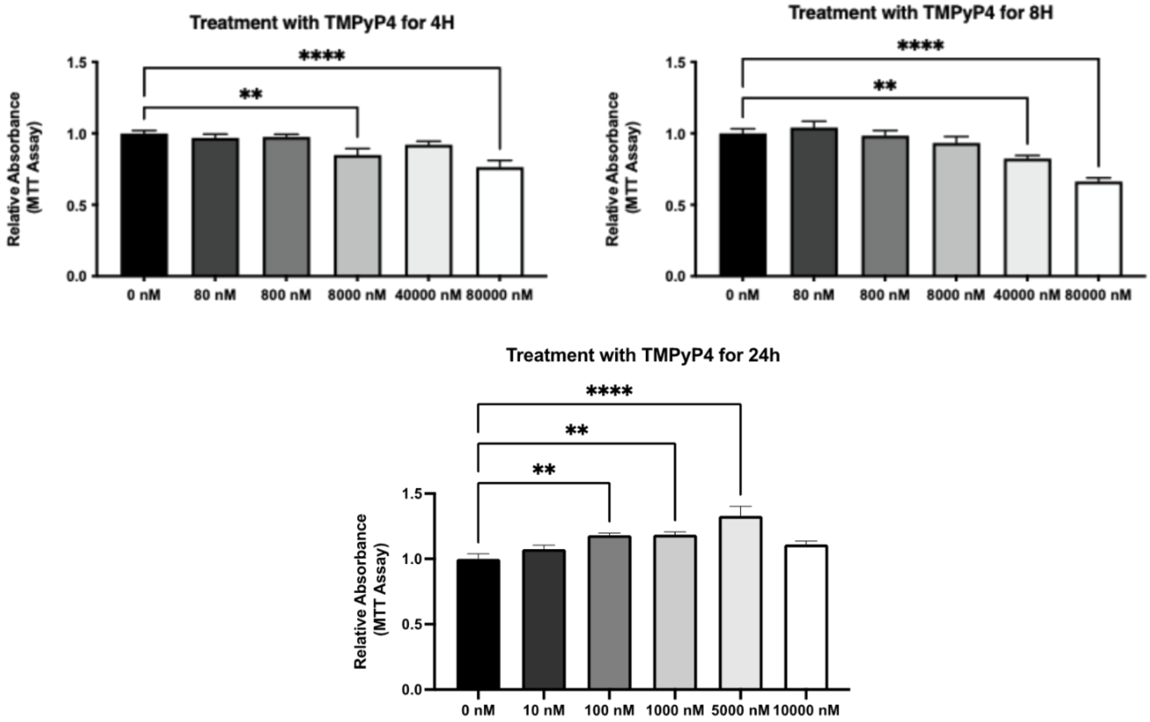


**Figure S9**. MTT assay shows the impact of TMPyP4 treatment on cell viability after 4, 8, and 24 hours at different dosages (as reported on the x-axis). Data are expressed as mean ± SEM of 8 replicate values in 3 independent experiments (n = 24, ***p<0.001, ****p<0.0001 vs 0nM).

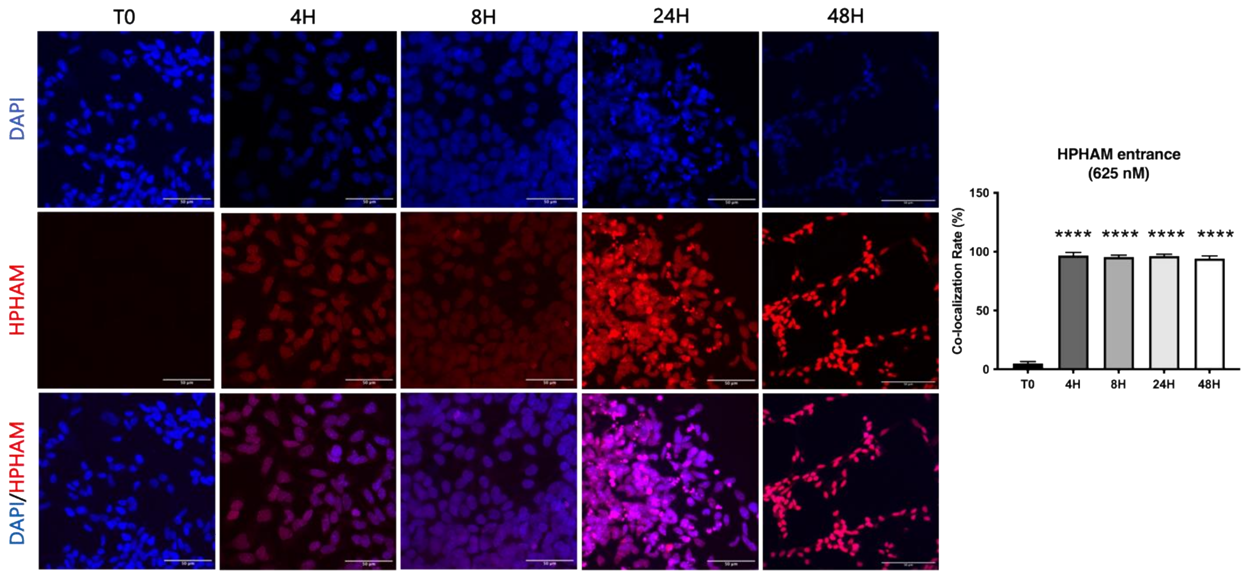


**Figure S10**. Immunofluorescence assay shows HPHAM entry in the cells after 0, 4, 8, 24, and 48 hours of treatment. HPHAM is shown in red, and nuclei are stained with DAPI in blue. Scale bar = 50 □m. Co-localization rate was calculated via Leica software and data is reported as a histogram on the right. (3 fields per experiment, 3 independent experiments; n = 9, ****p<0.0001 vs T0).

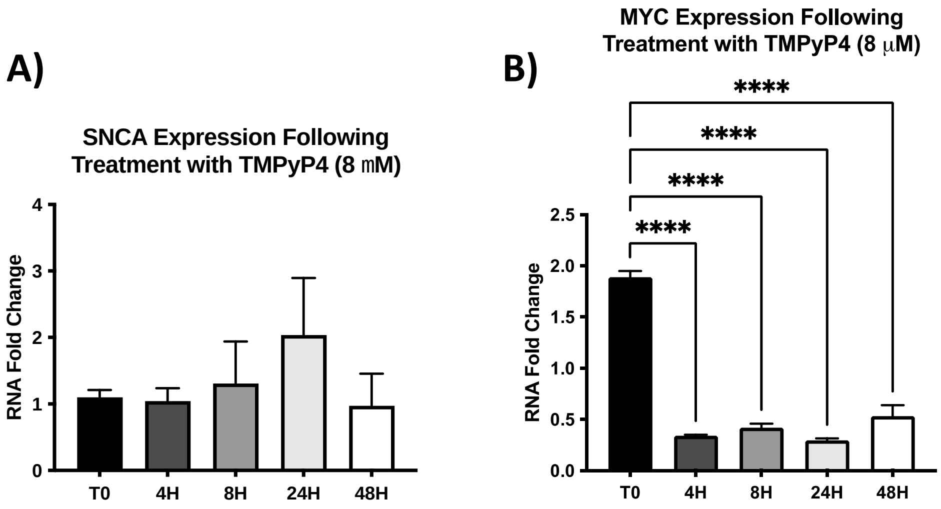


**Figure S11.** (A) SNCA and (B) MYC mRNA expression levels were assessed via Real-Time PCR following treatment with 8 μM TMPyP4 for 0, 4, 8, 24, and 48 hours. TBP was used as a housekeeping gene and data are expressed as mean ± SEM of 3 replicate values in 3 independent experiments (n=9; ****p<0.0001 vs T0).

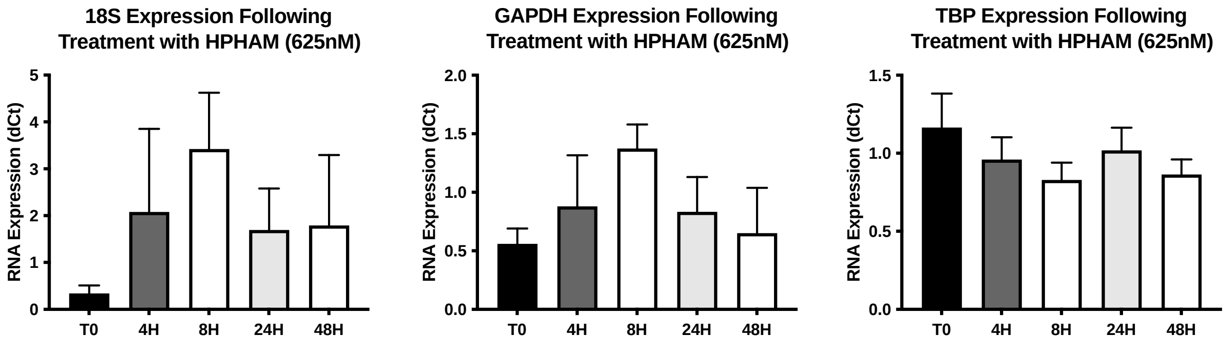


**Figure S12.** Real-Time PCR of different housekeeping genes shows that TBP is the most stable one following G4L treatment. 18S, GAPDH, and TBP mRNA expression levels were assessed via Real-Time PCR following treatment with 625nM HPHAM 0, 4, 8, 24, and 48 hours. Data are expressed as mean ± SEM of 3 replicate values in 1 independent experiment (n=3).

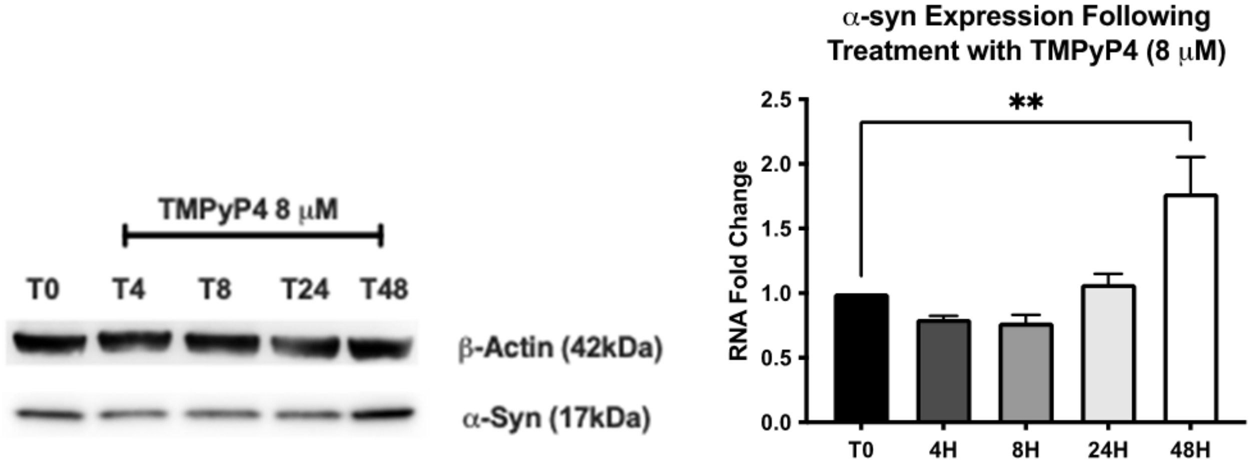


**Figure S13.** α-Syn expression levels were assessed via Western Blotting following treatment with 8 μM TMPyP4 for 0, 4, 8, 24, and 48 hours. β-Actin was used as a housekeeping protein and data are expressed as mean ± SEM of 1 experiment analyzed in triplicate (n=3; $^{**}p<0.01$ vs T0).

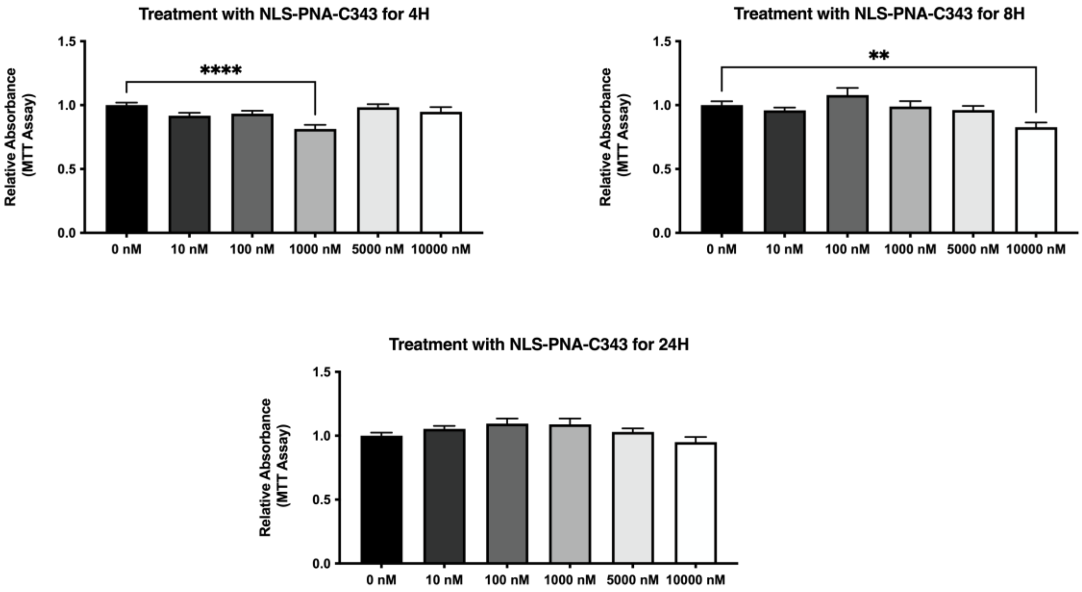


**Figure S14**. MTT assay shows the impact of NLS-PNA-C343 treatment on cell viability after 4, 8, and 24 hours at different dosages (as reported on the x-axis). Data are expressed as mean ± SEM of 8 replicate values in 3 independent experiments (n = 24, **p<0.01, ****p<0.0001 vs 0nM).

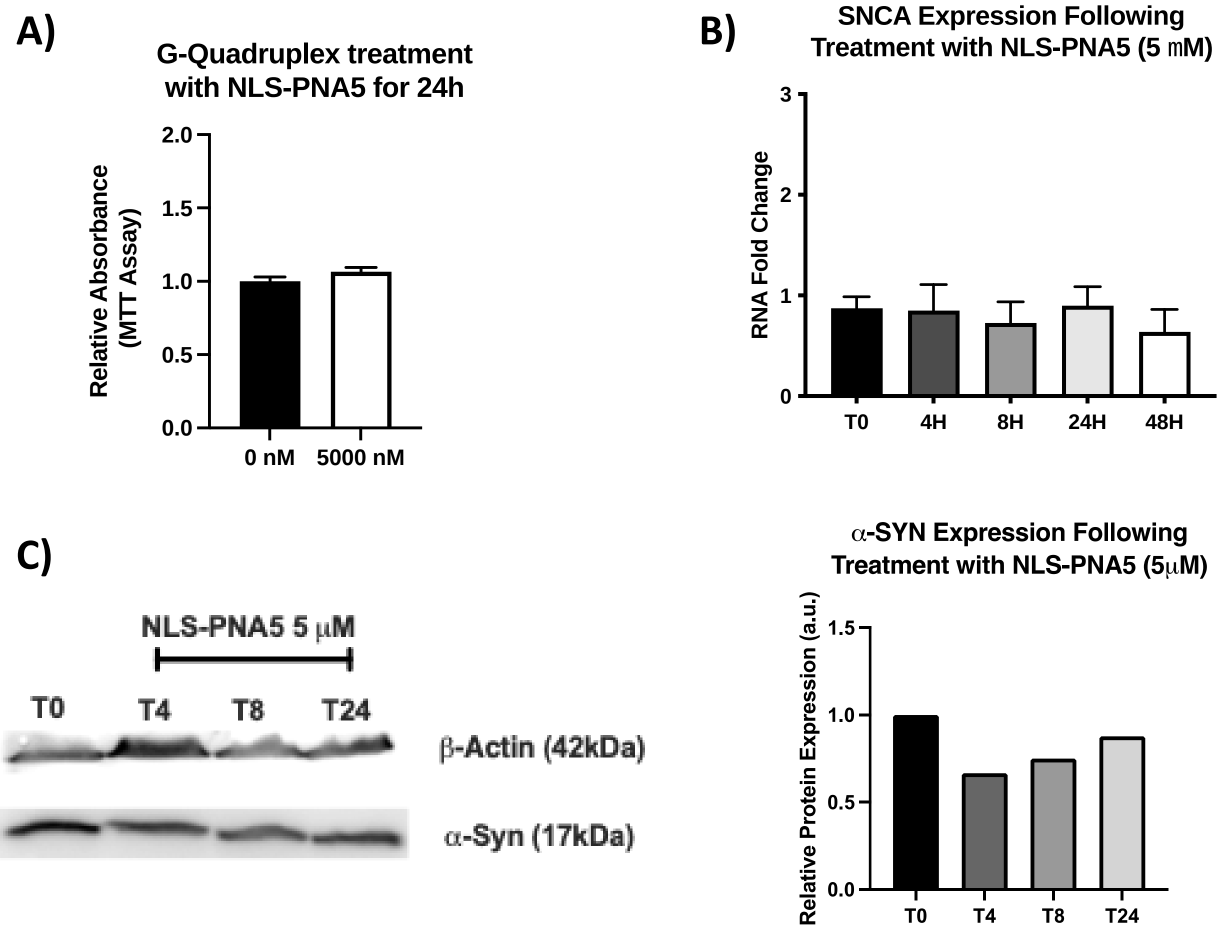

**Figure S15.** (A) MTT assay shows the impact of NLS-PNA5 treatment on cell viability after 24 hours at 5 μM. Data are expressed as mean ± SEM of 8 replicate values in 1 experiment (n = 8, p=ns). (B) SNCA mRNA expression levels were assessed via Real-Time PCR following treatment with 5 μM NLS-PNA5 for 0, 4, 8, 24, and 48 hours. TBP was used as a housekeeping gene and data are expressed as mean ± SEM of 3 replicate values in 1 experiment analyzed twice (n=6; p=ns). (C) α-Syn expression levels were assessed via Western Blotting following treatment with 5 μM NLS-PNA5 for 0, 4, 8, and 24 hours. β-Actin was used as a housekeeping protein and data are expressed as mean ± SEM of 2 independent experiments (n=2; p=ns).

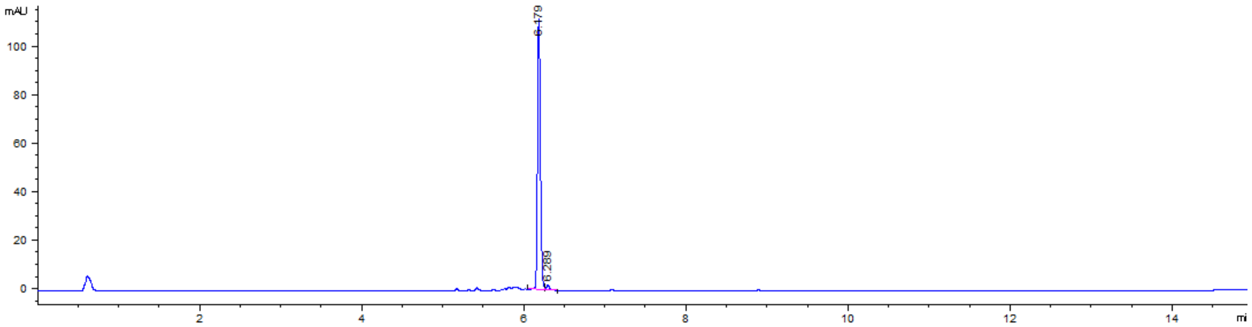


**Figure S16.** Analytic HPLC profile of HPHAM: r.T. 6.18 min; purity of 98.9%.

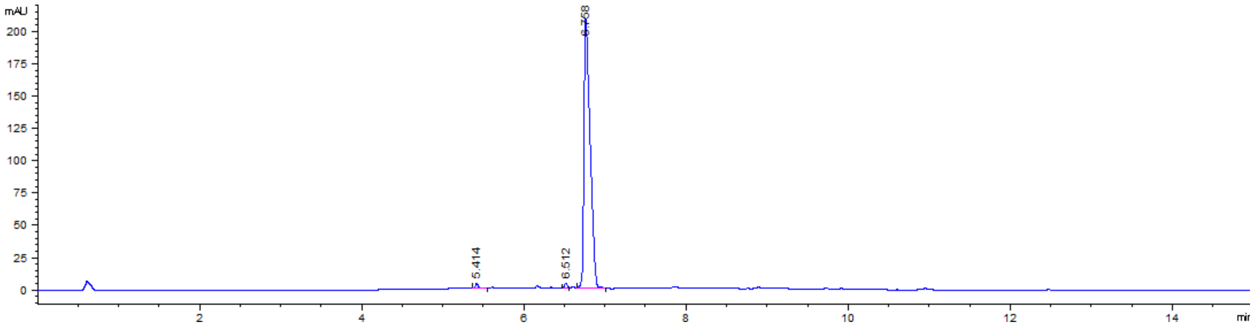


**Figure S17.** Analytic HPLC profile of PDS: r.T. 6.76 min; purity of 98.1%.

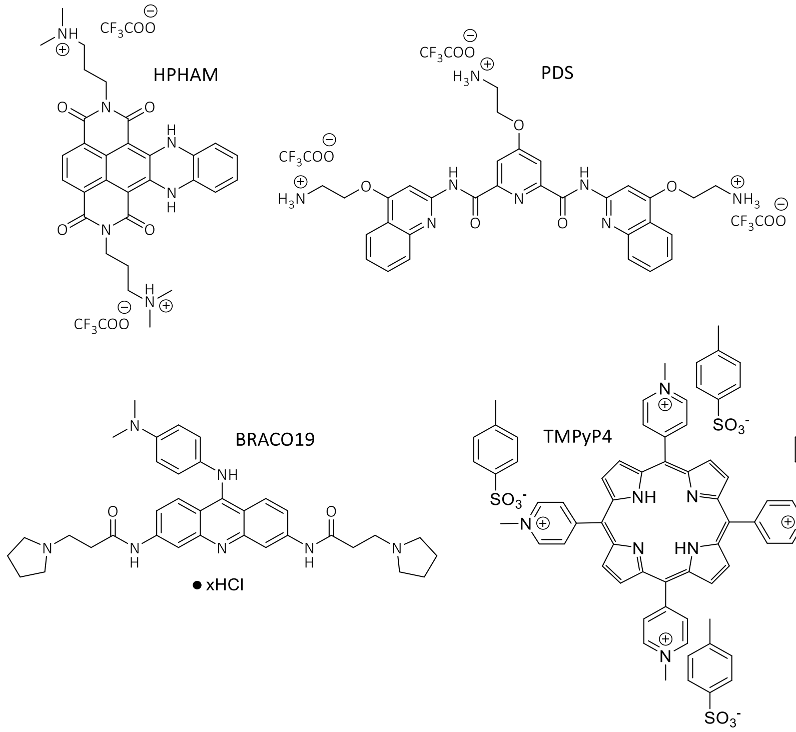


**Figure S18.** Chemical structures of the G4-ligands here used. HPHAM and PDS were synthesized in our laboratory, while Braco19 hydrochloride and TMPyP4 were purchased by Merck.

**Table S5.** Name and nucleobase sequences of the DNA and RNA model oligonucleotides here analyzed.

| **Name** | **Sequence (5'-3')** | **Length** | **GC content** |
|---|---|---|---|
| pSNCAex | GATGGGGATGGGGCAGGGGGCGCGGGGTGA | 30 nt | 76.7% |
| pSNCA | GGGGATGGGGCAGGGGGCGCGGGG | 24 nt | 87.5% |
| mSNCA | GGAGGAGGACUAGGAGGAGGAGG | 22 nt | 68.2% |
| FAM-pSNCAex-TAMRA | FAM-GATGGGGATGGGGCAGGGGGCGCGGGGTGA-TAMRA | 30 nt | 76.7% |
| FAM-pSNCA-TAMRA | FAM-GGGGATGGGGCAGGGGGCGCGGGG-TAMRA | 24 nt | 87.5% |
| FAM-mSNCA-TAMRA | FAM-GGAGGAGGACUAGGAGGAGGAGG-TAMRA | 22 nt | 68.2% |
| Myc-23456 | TGAGGGTGGGGAGGGTGGGGAAGG | 24 nt | 70.8% |
| cMyc | TGAGGGTGGGTAGGGTGGGTAA | 22 nt | 59.1% |
| 22ag | AGGGTTAGGGTTAGGGTTAGGG | 22 nt | 54.5% |
| CKIT-1 | AGGGAGGGCGCTGGGAGGAGGG | 22 nt | 77.3 % |
| LTR-III | GGGAGGCGTGGCCTGGGCGGGACTGGGG | 28 nt | 82.1% |

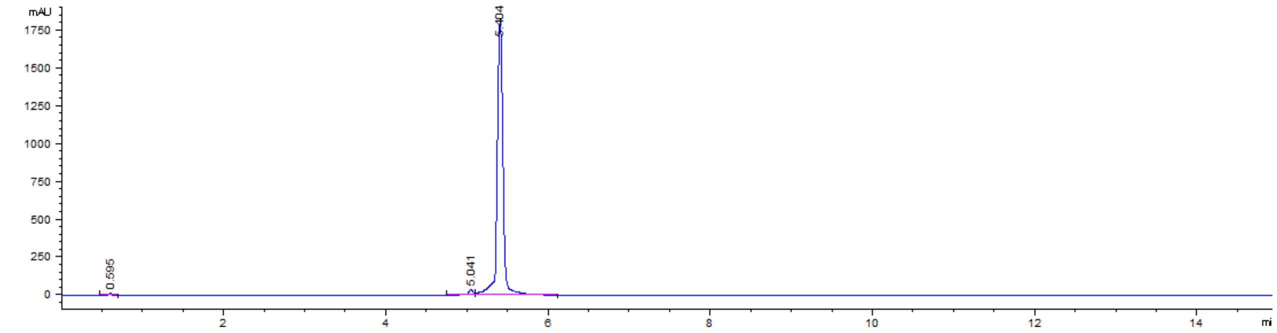


**Figure S19.** Analytic HPLC profile of NLS-PNA-C343: r.T. 5.40 min; purity of 97.6%.

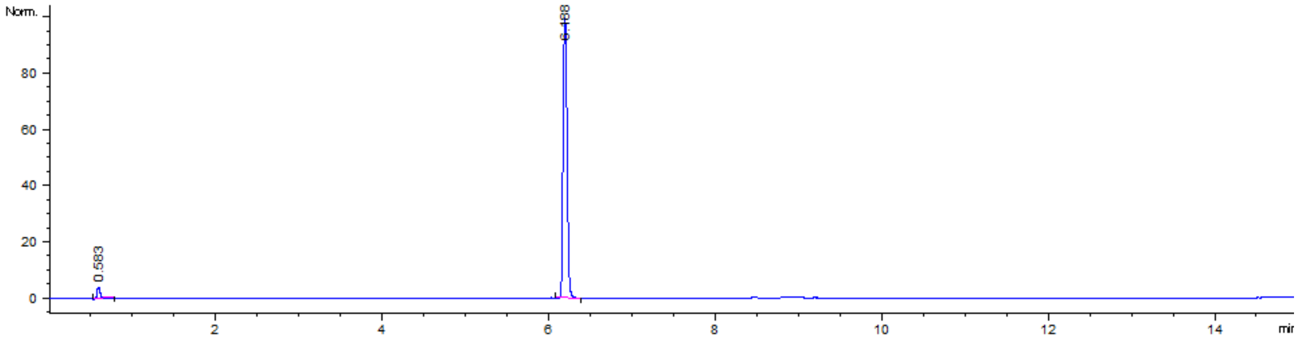


**Figure S20.** Analytic HPLC profile of NLS-PNA5: r.T. 6.19 min; purity of 98.5%.

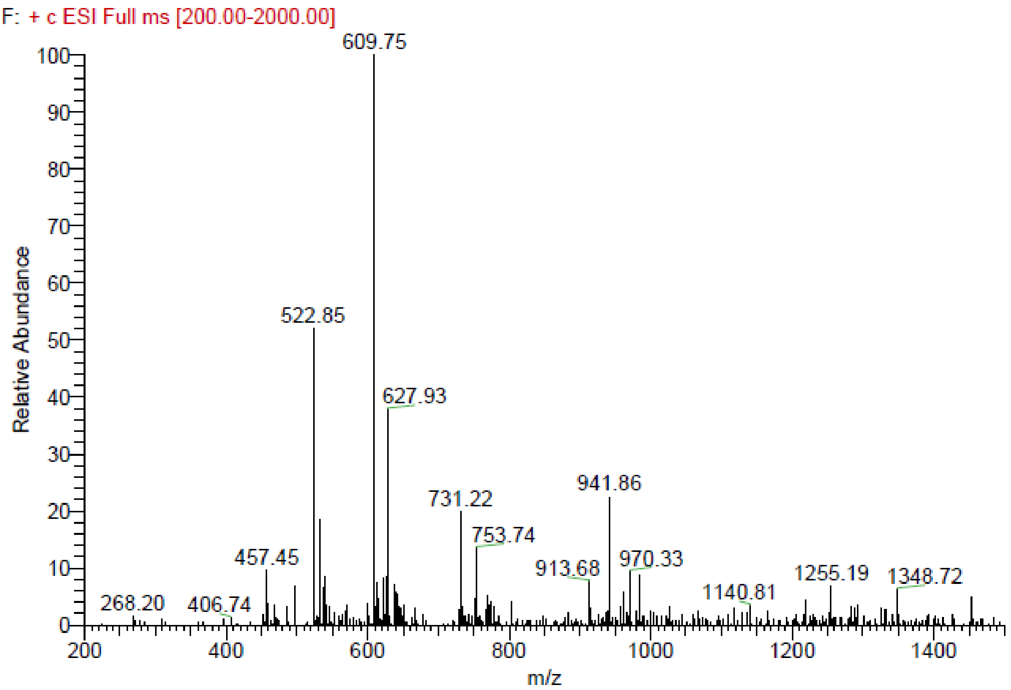


**Figure S21.** UHPLC-MS data of NLS-PNA-C343 (M.W.: 3651.9 u.m.a).

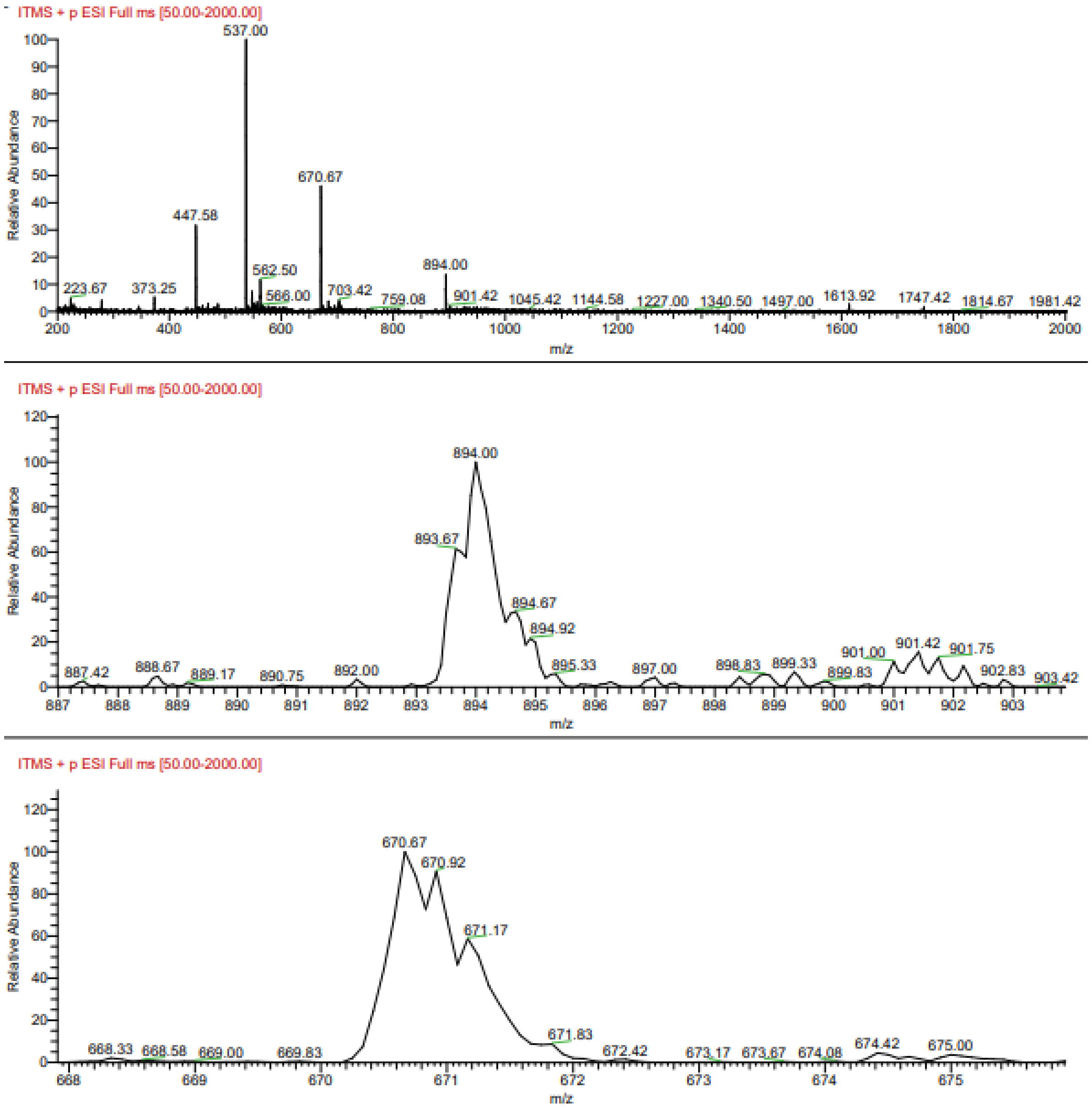


**Figure S22.** UHPLC-MS data of NLS-PNA5 (M.W.: 2677.6 u.m.a).

**Table S6.** List of primers used in this study.

| | | |
|---|---|---|
| NON_G4_GAPDH -FW | CCCCTTCATACCCTCACGTA | ChIP, RIP, qPCR Stop Assay |
| NON_G4_GAPDH-REV | GACAAGCTTCCCGTTCTCAG | ChIP, RIP, qPCR Stop Assay |
| SNCA_G4-FW | AACTTAACGTGAGGCGCAAA | ChIP, RIP, qPCR Stop Assay |
| SNCA_G4-REV | CTGATTTGTCAGCGCTTCTG | ChIP, RIP, qPCR Stop Assay |
| G4_cMYC-FW | CATGCGGCTCTCTTACTCTG | ChIP, RIP, qPCR Stop Assay |
| G4_cMYC-REV | CGGAGATTAGCGAGAGAGGA | ChIP, RIP, qPCR Stop Assay |
| SNCA-FW | TGCTGCTGAGAAAACCAAAC | Real-Time PCR |
| SNCA-REV | GAAGCACCGAAATGCTGAGT | Real-Time PCR |
| TBP-FW | TTAACTTCGCTTCCGCTGGC | Real-Time PCR |
| TBP-REV | CAAGAAACAGTGATGCTGGGTCA | Real-Time PCR |
| MYC-FW | CTGGGAAGAAGCCAGTTCAG | Real-Time PCR |
| MYC-REV | TGGGCCATAGGTTTTCAGAG | Real-Time PCR |
| GAPDH-FW | AAAGTGAAGGTCGGAGTACAACGGATTTGGT | Real-Time PCR |
| GAPDH-REV | AGCCTTGACGGTGCCATGGAATTTGCCATG | Real-Time PCR |
| 18S-FW | AGTACGCACGGCCGGTACAGTGAAACTGCG | Real-Time PCR |
| 18S-REV | CGGGTTGGTTTTGATCTGATAAATGCACGC | Real-Time PCR |

**Denaturation curves obtained by CD-melting experiments.**

Denaturation curves obtained for 2.5 µM pSNCA and pSNCAext in 10 mM Lithium cacodylate buffer pH 7.4, with different KCl (from 100 mM to 1 mM)/LiCl (from 0 mM to 99 mM) amounts.

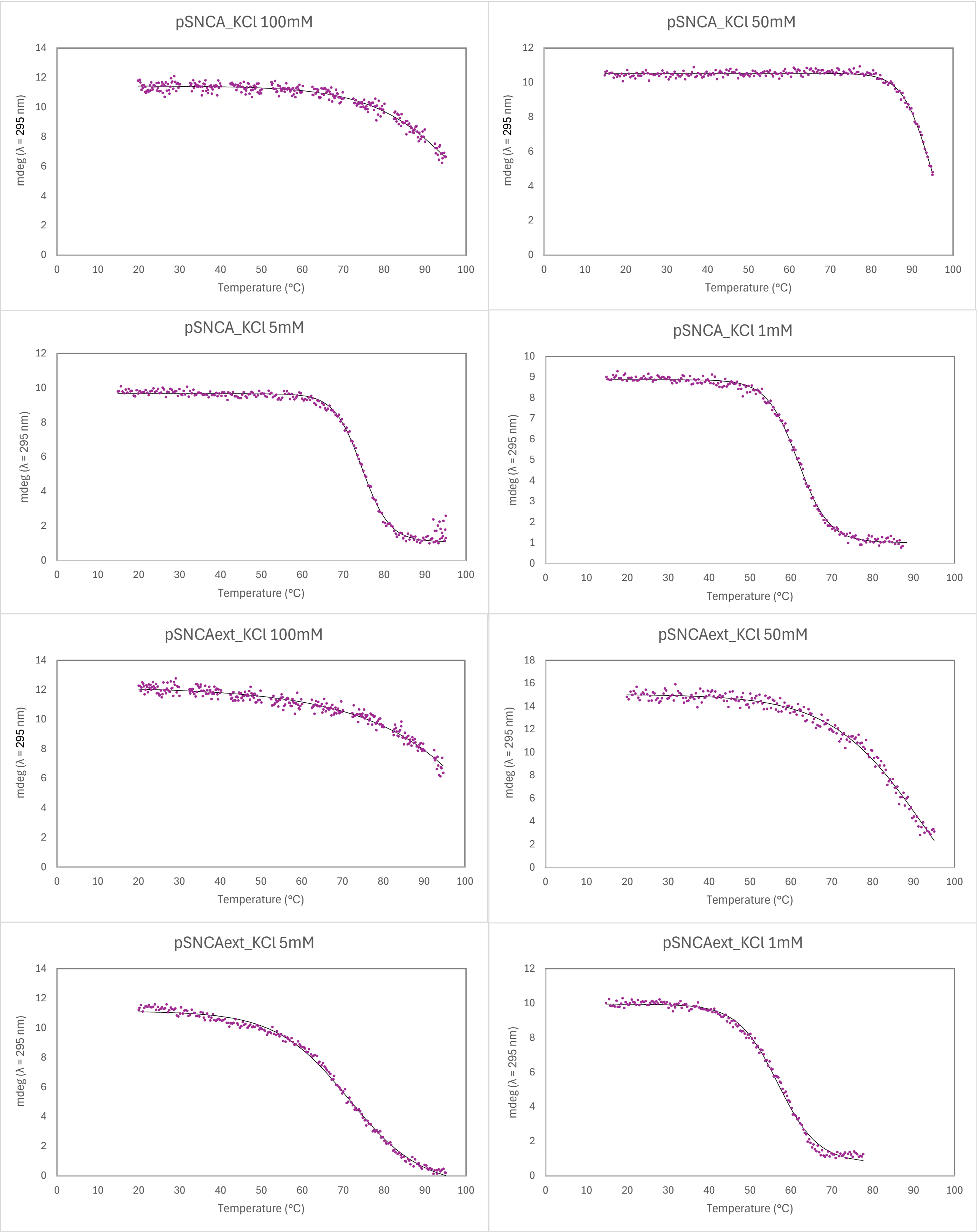

Denaturation curves obtained for 2.5 μM pSNCA and pSNCAext in the presence of 4 equivalent stabilizing ligands in 10 mM Lithium cacodylate buffer pH 7.4, with 1 mM KCl and 99 mM LiCl.

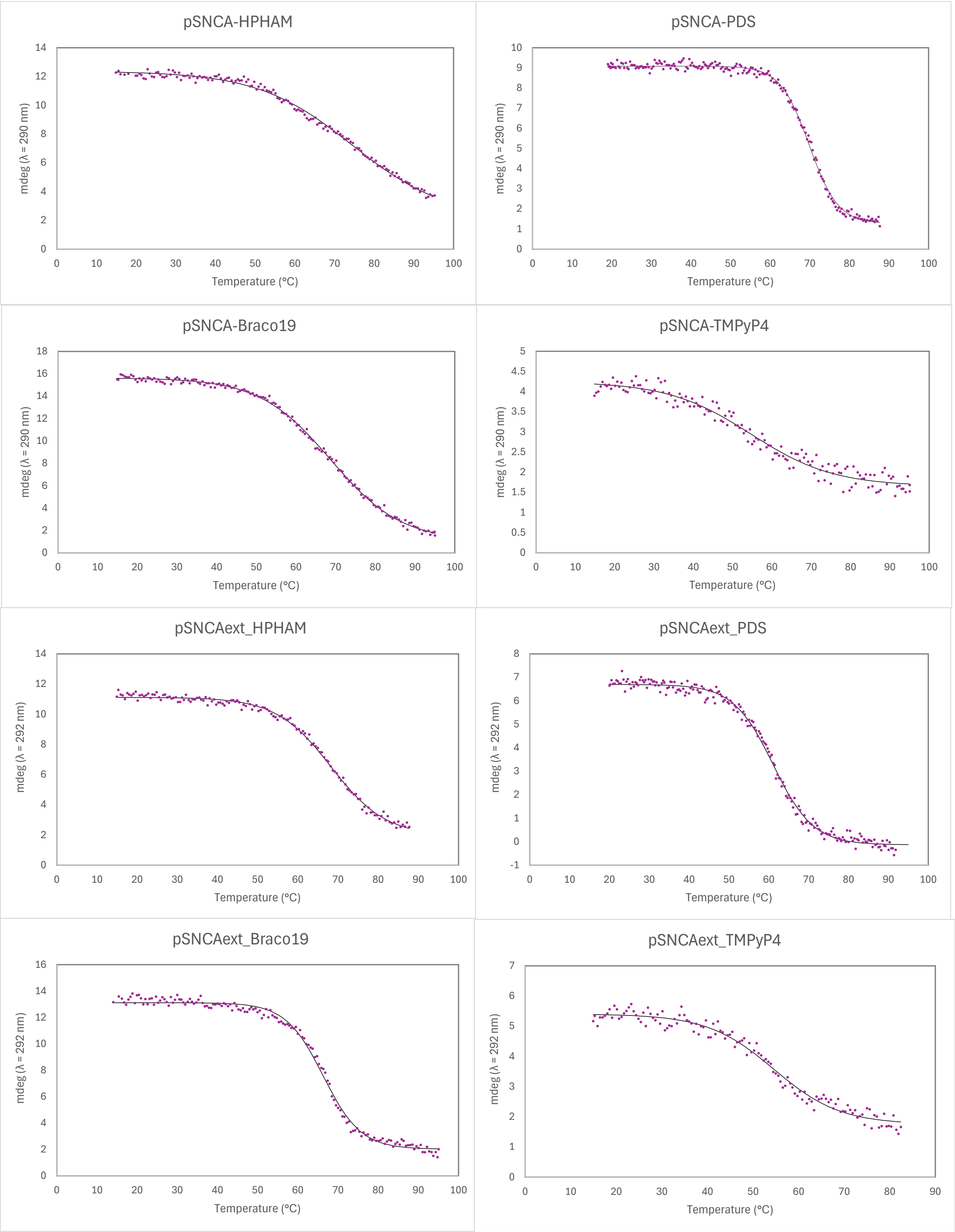

Denaturation curves obtained for 2.5 µM pSNCAext in the presence of 4 equivalent NLS-PNA-C343 or NLS-PNA5 in 10 mM Lithium cacodylate buffer pH 7.4, with 100 mM KCl.

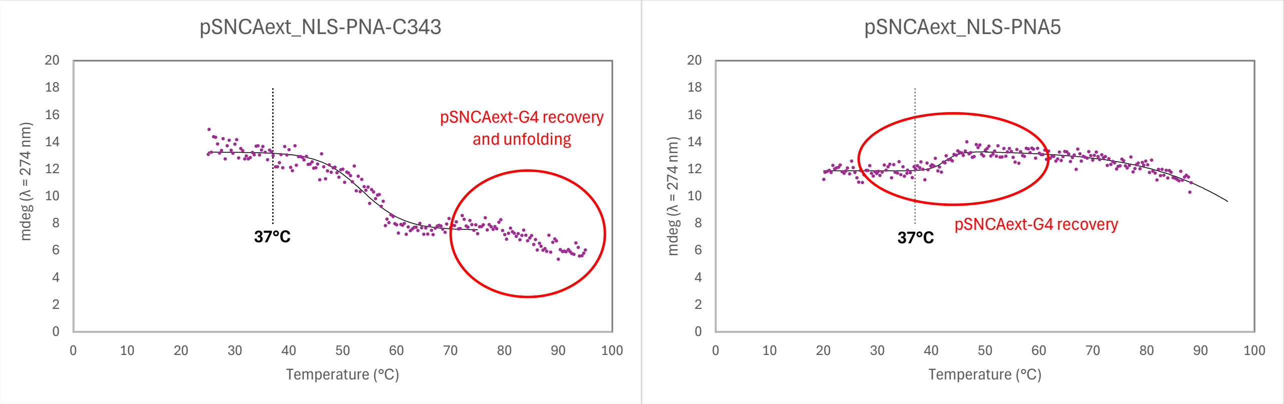


Denaturation curves obtained for 2.5 µM mSNCA with 10 mM or 100 mM KCl in 10 mM Lithium cacodylate buffer pH 7.4.

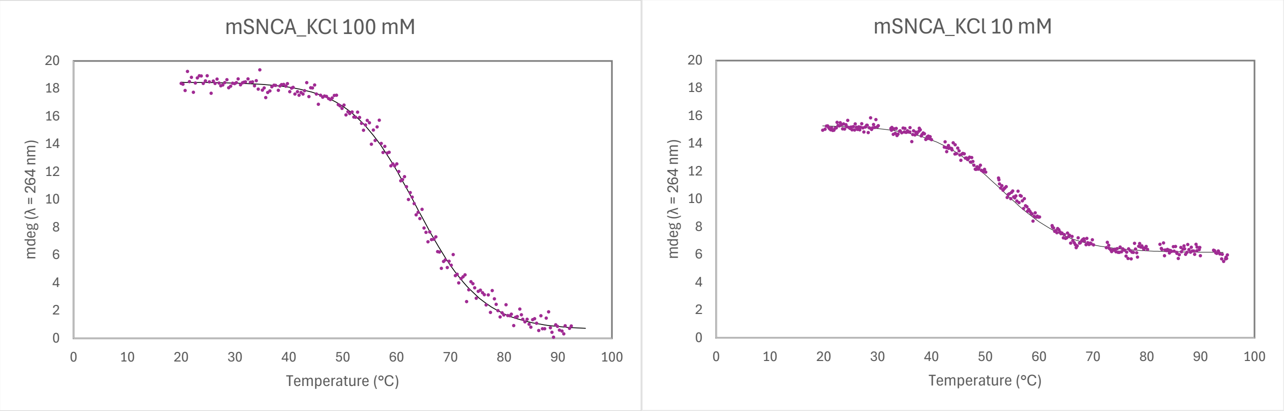

Denaturation curves obtained for 2.5 µM mSNCA in the presence of 4 equivalent stabilizing ligands with 100 mM KCl in 10 mM Lithium cacodylate buffer pH 7.4.

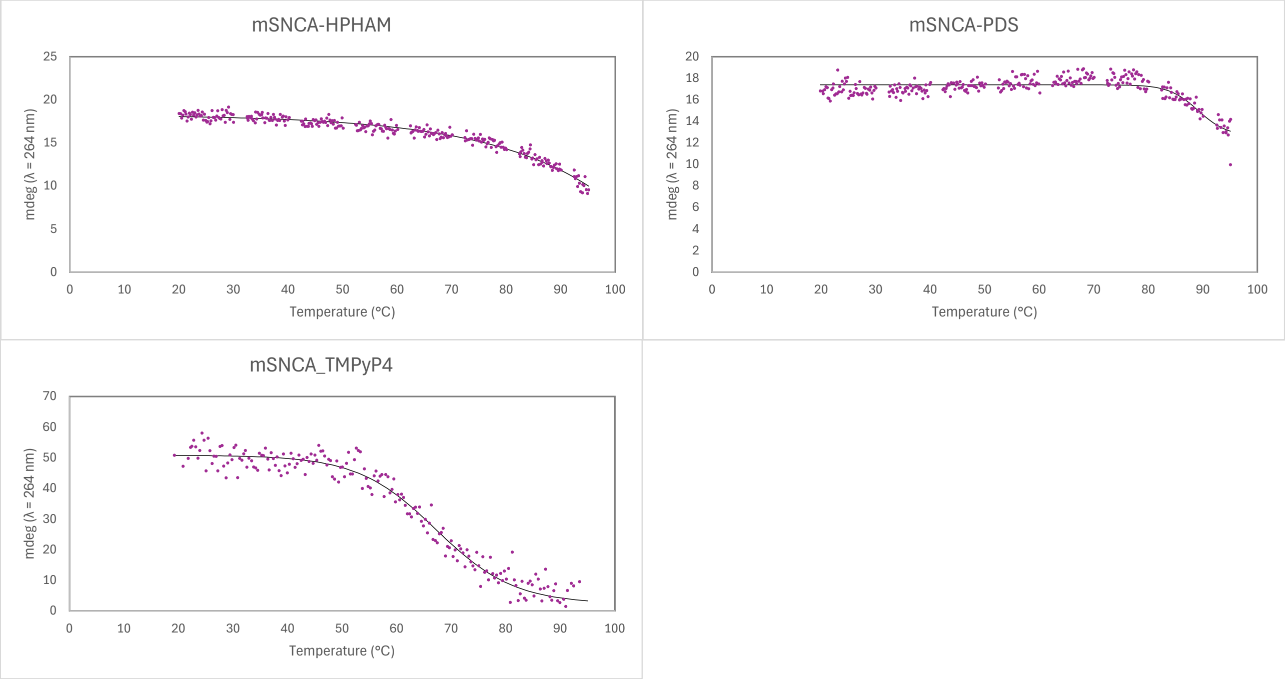

Denaturation curves obtained for 2.5 µM Myc23456, cMyc, 22ag, LTRIII, cKIT-1 in 10 mM Lithium cacodylate buffer pH 7.4, with different KCl/LiCl concentrations.

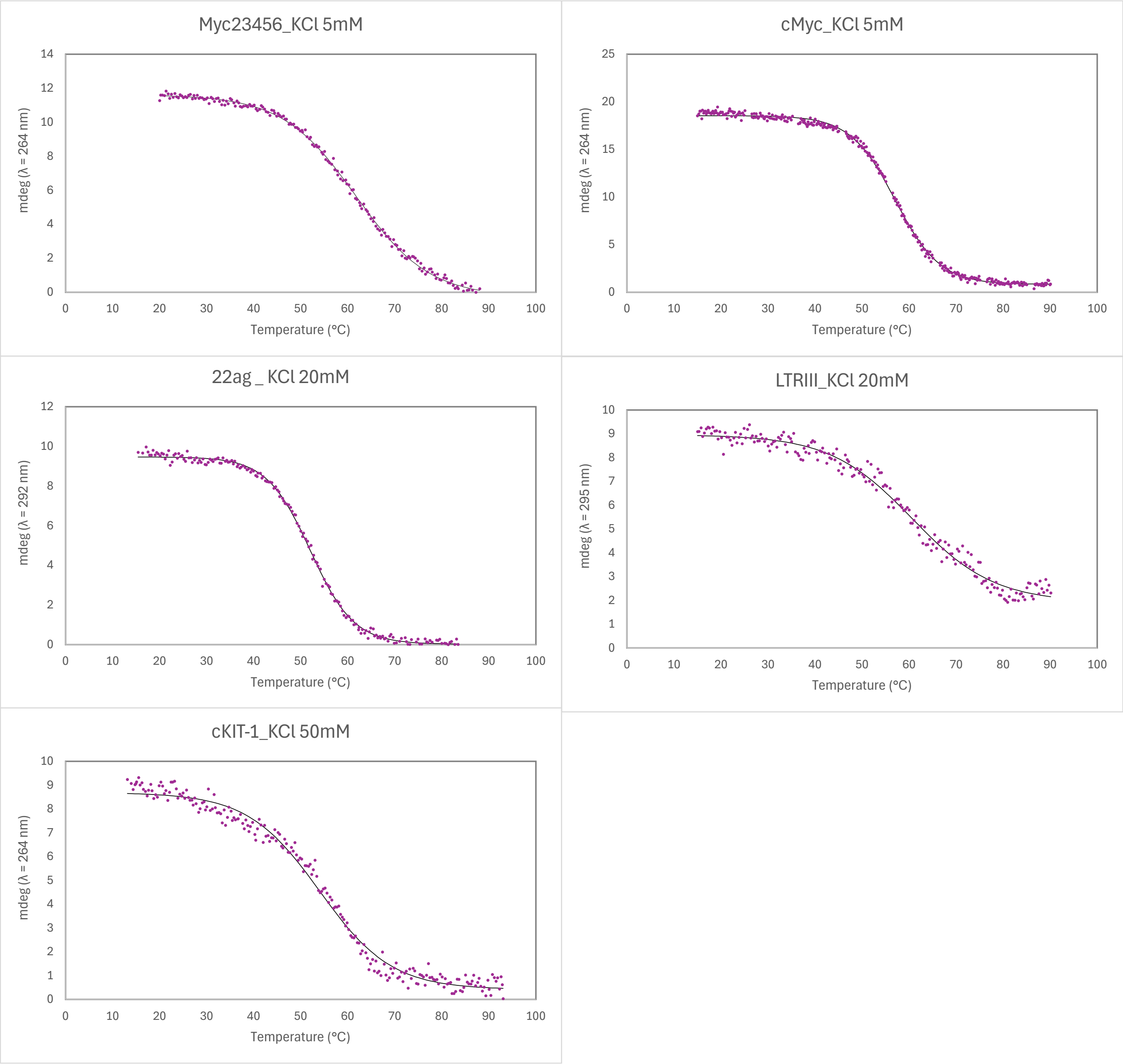

**Denaturation curves obtained by UV-melting experiments.**

Denaturation curves obtained for 2.5 µM pSNCA, pSNCAex, and mSNCAt in 10 mM Lithium cacodylate buffer pH 7.4, with different KCl /LiCl concentrations.

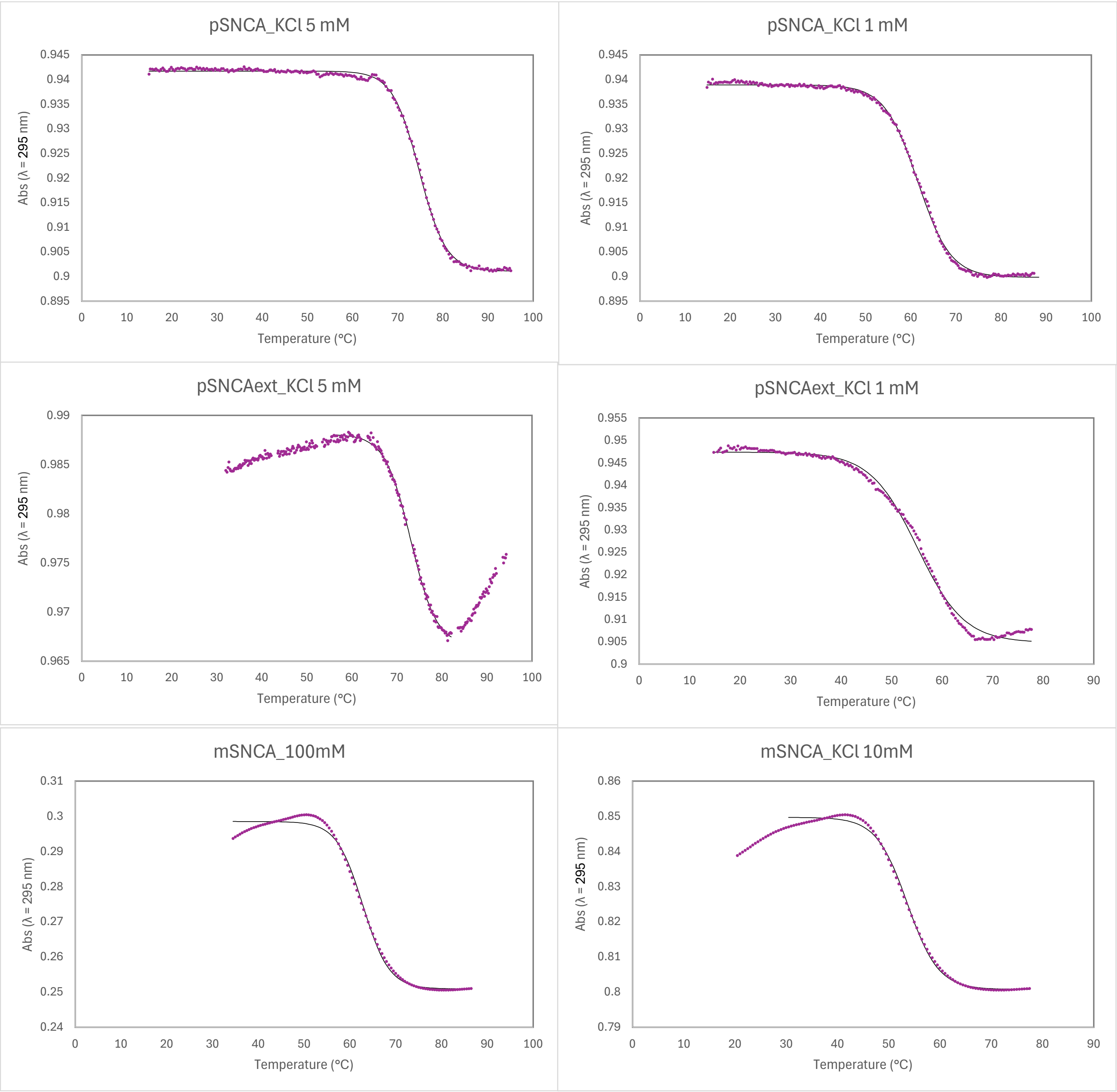

**Denaturation curves obtained by FRET-melting experiments.**

Denaturation curves obtained for 0.25 µM pSNCA, pSNCAext, and mSNCA in 10 mM Lithium cacodylate buffer pH 7.4, with different KCl amounts.

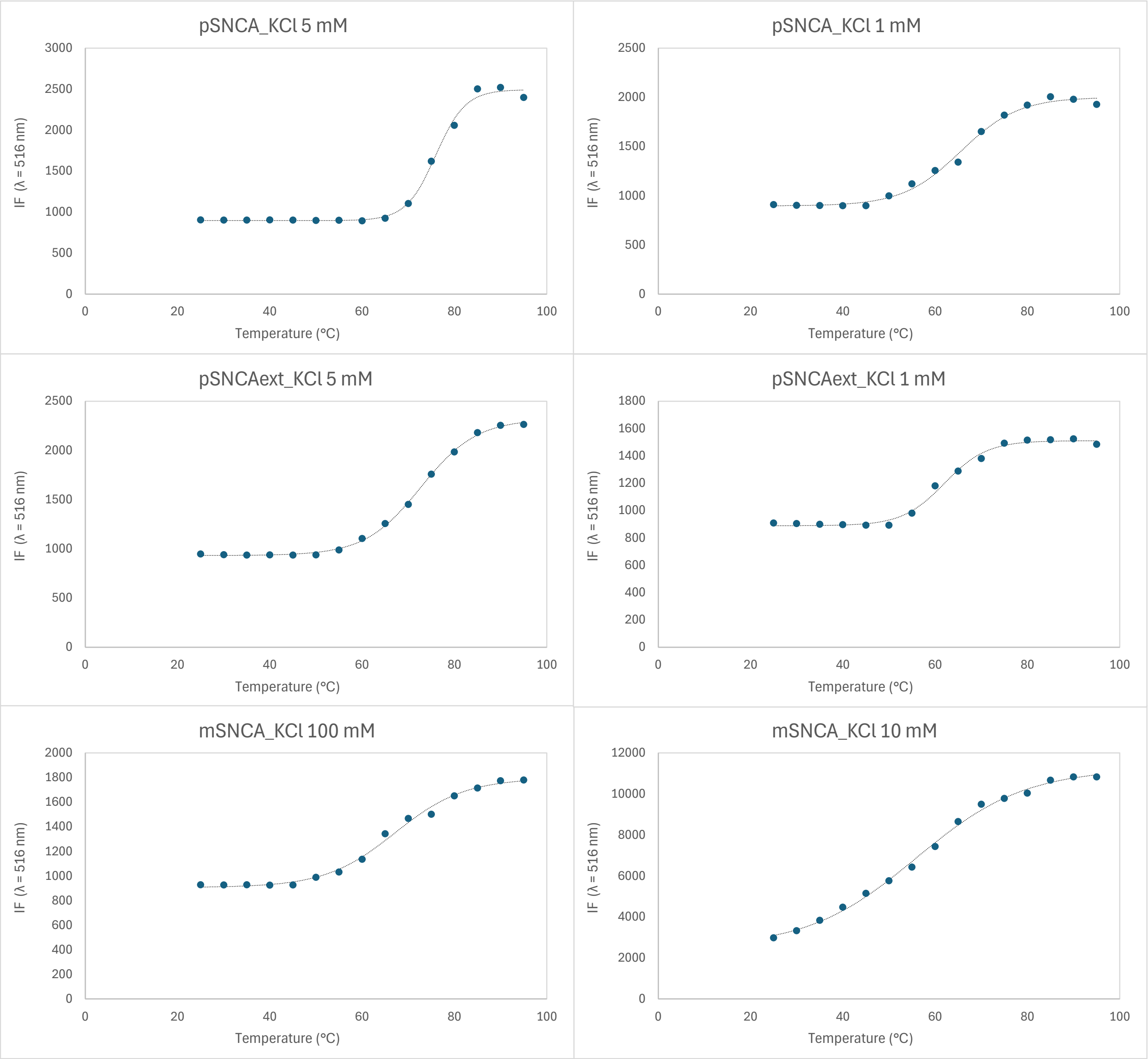

Denaturation curves obtained for 0.25 µM pSNCA and pSNCAext in the presence of 4 equivalent stabilizing ligands in 10 mM Lithium cacodylate buffer pH 7.4, with 1 mM KCl and 99 mM LiCl.

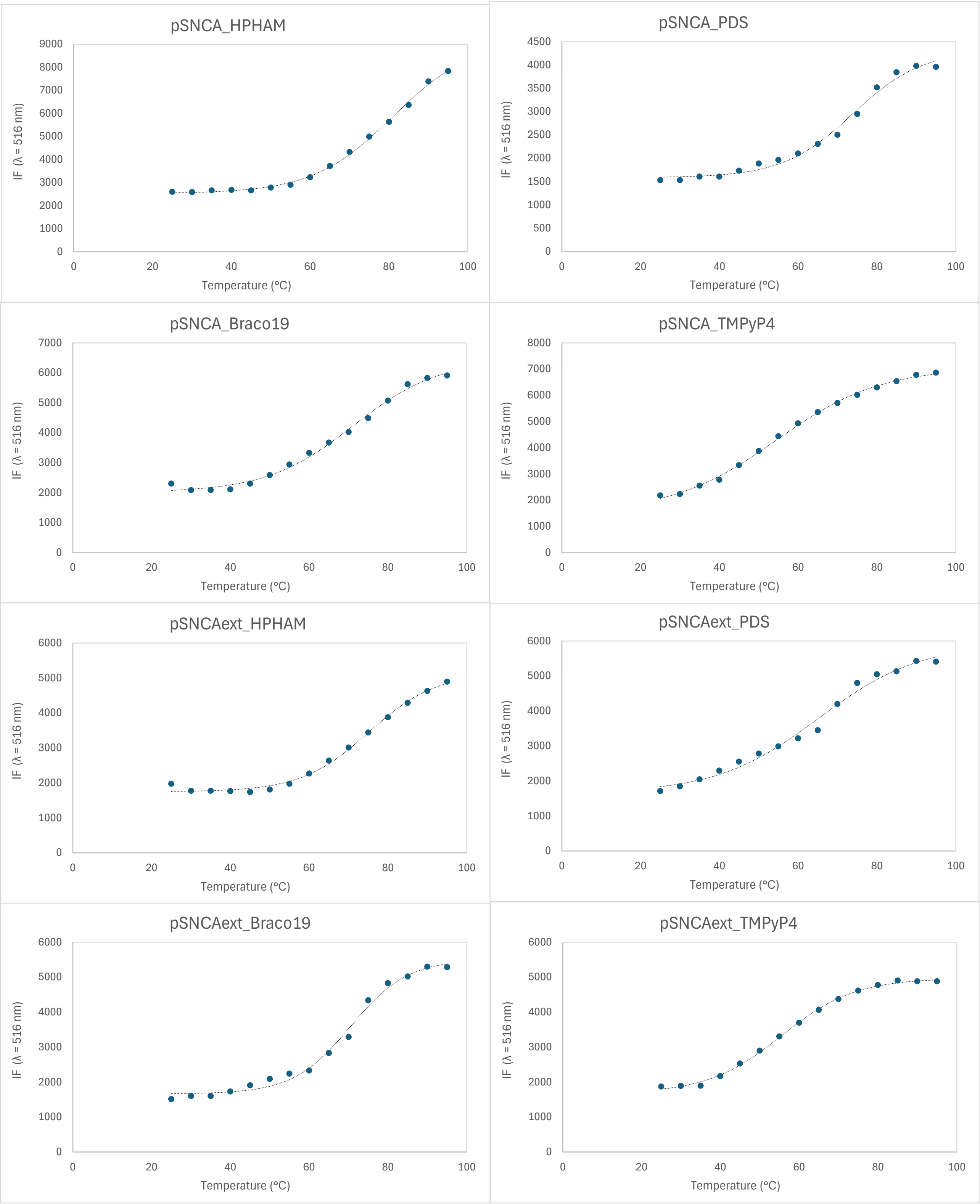

Denaturation curves obtained for 0.25 μM mSNCA in the presence of 4 equivalent of HPHAM, PDS, or TMPyP4 in 10 mM Lithium cacodylate buffer pH 7.4, with 100 mM KCl.

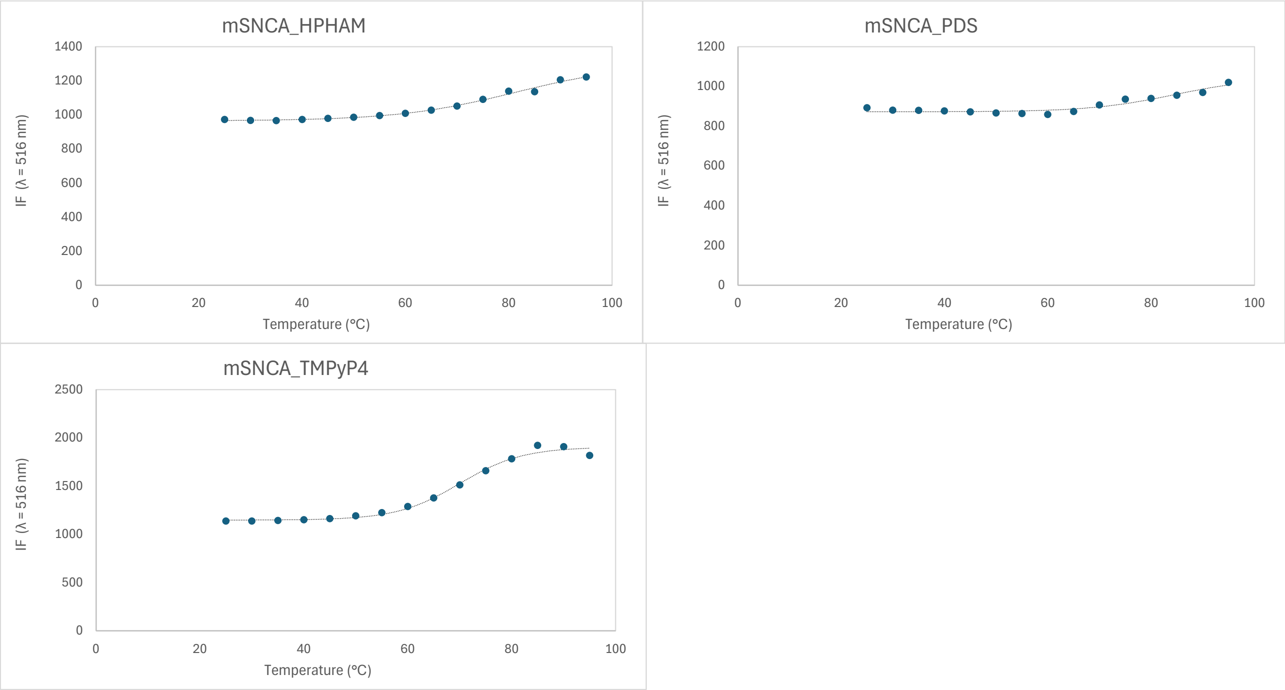
